# Population genomics and demographic sampling of the ant-plant *Vachellia drepanolobium* and its symbiotic ants from sites across its range in East Africa

**DOI:** 10.1101/475392

**Authors:** John H. Boyle, Dino Martins, Paul M. Musili, Naomi E. Pierce

## Abstract

The association between the African ant plant, *Vachellia drepanolobium*, and the ants that inhabit it has provided insight into the boundaries between mutualism and parasitism, the response of symbioses to environmental perturbations, and the ecology of species coexistence. We use a landscape genomics approach at sites sampled throughout the range of this system in Kenya to investigate the demographics and genetic structure of the different partners in the association. We find that different species of ant associates of *V. drepanolobium* show striking differences in their spatial distribution throughout Kenya, and these differences are only partly correlated with abiotic factors. A comparison of the population structure of the host plant and its three obligately arboreal ant symbionts, *Crematogaster mimosae*, *Crematogaster nigriceps*, and *Tetraponera penzigi*, shows that the ants exhibit somewhat similar patterns of structure throughout each of their respective ranges, but that this does not correlate in any clear way with the respective genetic structure of the populations of their host plants. A lack of evidence for local coadaptation in this system suggests that all partners have evolved to cope with a wide variety of biotic and abiotic conditions.

## Introduction

The Whistling Thorn acacia, *Vachellia drepanolobium* (formally referred to as *Acacia drepanolobium*), and its several species of resident ants comprise a well-studied symbiosis that dominates savannahs throughout its range in Eastern Africa. The ants nest only in hollow, swollen-thorn domatia of this *Vachellia* species, and receive extrafloral nectar from the host tree. In return, the ants typically attack other organisms that venture onto the trees, thereby deterring herbivores and potential enemies (Hocking 1970, Madden and Young 1992). Because ants of different species compete to occupy *V. drepanolobium* domatia, the system has been used to investigate possible mechanisms of species coexistence (e.g., Stanton *et al.* 2002, Palmer 2004, Boyle *et al.* 2017). At a well-studied field site in Laikipia, Kenya, experiments have shown that associated ant species vary in the benefits they provide to the tree as well as the costs they impose, and analyses of their interactions with their host plants at that site have yielded insights into the boundaries between parasitism and mutualism (Palmer *et al.* 2010, Martins 2010, Stanton and Palmer 2011). Because of the considerable human impacts on the savannah, much work has focused on elucidating how the system as a whole responds to environmental perturbation (Palmer *et al.* 2008, Riginos *et al.* 2015).

The full range of *V. drepanolobium* roughly spans from Tanzania in the south, Uganda in the west and southern Ethiopia in the North, although it is most abundant across the savannahs of Tanzania and Kenya, where it is restricted to black cotton soils. This study examines *V. drepanolobium* and its ant associates at locations distributed throughout Kenya. At each site, we describe both sides of the association, recording tree sizes and densities, as well as which ants are present at each site and in what numbers. The sites encompass a geographically diverse area: two are in Laikipia in north central Kenya, three are in the highlands area south of Nairobi, three are in the Rift Valley, and two sites lie on the edges (one western and one eastern) of the Rift. Together with this demographic survey, we report the results of a population genomic study of *V. drepanolobium* and its three obligate phytoecious (i.e., tree-dwelling) ant inhabitants, *Crematogaster mimosae*, *C. nigriceps*, and *Tetraponera penzigi*.

We first asked whether gene flow among populations of the host tree show patterns that differ from those found among its respective symbiotic ant species? The dynamics of both partners have been assessed in relatively few symbiotic systems, primarily macrobe-microbe interactions. In such symbioses, a wide range of possible outcomes have been described: population structures may be broadly congruent, as in a fungus-beetle symbiosis (Roe *et al.* 2011) or some lichen (Werth and Scheidegger 2012, Widmer *et al.*2012). Such congruence may signal specialized coevolution and reciprocal adaptation of populations of hosts and their respective symbionts, or can have other explanations, such as the apparent pattern of codiversification generated by shared vicariance events (discussed in Segraves 2010). On the other hand, some population studies of symbiosis reveal distinct population structures between host and symbiont, as in a cnidarian-dinoflagellate mutualism (Thornhill *et al.* 2013), a lichen host that showed less population structure than its algal symbiont (Werth and Sork 2010), or in the leaf-cutter ant-fungus interaction, a situation where many host ant species associate with a single fungal symbiont (Mikheyev *et al.* 2006). In contrast, in symbioses between two macroorganisms, population structures are broadly congruent, as in the ant-plant systems *Leonardoxa africana* (Leotard *et al.* 2009) or *Barteria fistulosa* (Blatrix *et al.* 2017), although the landscape genetics of symbioses between macroorganisms have been studied in relatively few cases, primarily addressing the population genetics of just one of the participants (e.g. Quek *et al.* 2007). We thus do not know whether the congruence in population structure that has so far been observed is a general feature of macrosymbioses.

A population genetic comparison of partners in the *V. drepanolobium* symbiosis is particularly interesting because of the well-documented behavioral variation in the ant partners. At a landscape scale, do we see signs of the same behaviors observed at Mpala, particularly the hierarchy in short-range colonization ability? Do the different partners show evidence of local adaption to regional environments, both abiotic (i.e., temperature, rainfall), and biotic (i.e., the presence or absence of certain species or genotypes)?

Answering these questions would enable us to extend the substantial body of research that has been done at sites in Laikipia, Kenya to sites throughout the range of *V. drepanolobium*. These questions are also important for understanding symbiosis more generally, especially non-pairwise, multi-partner symbioses. Symbioses between species often involve the asymmetrical exchange of different goods and services, and are typically not geographically homogenous (Stanton 2003, Thompson 2005). However, our understanding of the consequences of this complexity on population structure and gene flow is still lagging. Here we address this question by comparing the population structures of *V. drepanolobium* and its three common, obligate ant associates.

## Materials and Methods

### Site survey

Between 2013-2016, we surveyed *V. drepanolobium* and its ant associates at 10 sites in Kenya (Table 1, Figure 1). These sites included five in the highlands east of the Great Rift Valley and five in or around the Rift itself. The Rift Valley is a central feature of Kenyan geography. The Eastern Rift Valley runs through the middle of Kenya, from Lake Turkana in the north to the Tanzanian border in the south. Its formation began approximately 20 million years ago as the African tectonic plate began to split apart, a process that continues to this day (Chorowicz 2005). The difference between the relatively low-lying sites in the Rift and the highlands sites outside the Rift can be as much as a kilometer (Table 1), and the valley is separated from the highlands by a steep escarpment. The Rift Valley thus may be a barrier to gene flow in organisms including plants (Omondi *et al.* 2010, Ruiz Guajardo *et al.* 2010), insects (Lehmann *et al.* 1999) and vertebrates (Arctander *et al*. 1999, Ahlering *et al.* 2012).

**Table 1:** Demography of *V. drepanolobium*-ant association varies throughout Kenya

| Site | Cluster | Lat. (°) | Long. (°) | Alt. (m) | Plot size (m<sup>2</sup>) | Trees surveyed | Tree density (trees/ha) | Tree height (m) |
| --- | --- | --- | --- | --- | --- | --- | --- | --- |
| 1. Suyian Ranch | NE | 0.50675 | 36.69235 | 1900 | 8000 | 533 | 666 | 1.20 ± 0.36 |
| 2. Mpala Research Center | NE | 0.28961 | 36.87106 | 1807 | 67200<sup>2</sup> | 3391<sup>2</sup> | 505 | 2.41 ± 1.03 |
| 3. Kitengela | SE | -1.39645 | 36.82562 | 1673 | 2000 | 163 | 815 | 1.30 ± 0.52 |
|  | SE | -1.47344 | 36.64562 | 1947 |  |  |  |  |
| 4. Ngong Hills<sup>3</sup> |  | -1.47590 | 36.64340 | 1921 | 2000 | 138 | 690 | 2.67 ± 1.81 |
|  | SE | -1.60041 | 36.75144 | 1811 |  |  |  |  |
| 5. Isinya Road<sup>3</sup> |  | -1.60114 | 36.75115 | 1801 | 1500 | 163 | 1090 | 1.41 ± 0.52 |
| 6. Mogotio | NW | 0.17323 | 36.08356 | 1238 | 1040 | 44 | 420 | 4 ± 2 |
| 7. Koriema | NW | 0.47942 | 35.85226 | 1375 | 1050 | 28 | 270 | 5 ± 2 |
| 9. Gilgil | SW | -0.47670 | 36.35554 | 1938 | 1350 | 30 | 220 | 1.54 ± 0.40 |
| 10. Lake Naivasha | SW | -0.82773 | 36.33415 | 1195 | 2000 | 76 | 380 | 1.65 ± 0.57 |
| 11. Narok Road | SW | -1.10153 | 36.06934 | 2073 | 2250 | 183 | 813 | 0.87 ± 0.17 |

Proportion of trees occupied by:
| Site | Cluster | *C. sjostedti* | *C. mimosae* | *C. nigriceps* | *T. penzigi* | None<sup>1</sup> | Other<sup>1</sup> |
| --- | --- | --- | --- | --- | --- | --- | --- |
| 1. Suyian Ranch | NE | 0.48 | 0.19 | 0.21 | 0.08 | 0.02 | 0.01 |
| 2. Mpala R.C. | NE | 0.21 | 0.49 | 0.14 | 0.11 | 0.04 | 0.02 |
| 3. Kitengela | SE | 0 | 0.14 | 0.67 | 0.13 | 0.03 | 0.02 |
| 4. Ngong Hills<sup>3</sup> | SE | 0 | 0.73 | 0.11 | 0.09 | 0.07 | 0 |
| 5. Isinya Road<sup>3</sup> | SE | 0 | 0.16 | 0.74 | 0.09 | 0 | 0.01 |
| 6. Mogotio | NW | 0 | 0 | 0<sup>4</sup> | 0.05 | 0 | 0.95<sup>4</sup> |
| 7. Koriema | NW | 0 | 0 | 0.57 | 0.39 | 0.04 | 0 |
| 9. Gilgil | SW | 0 | 0 | 0.03 | 0.20 | 0.77 | 0 |
| 10. Lake Naivasha | SW | 0 | 0<sup>5</sup> | 0<sup>5</sup> | 0.84 | 0.16 | 0 |
| 11. Narok Road | SW | 0 | 0.11 | 0.69 | 0.04 | 0.16 | 0 |

Average height (m) of trees occupied by:
| Site | Cluster | *C. sjostedti* | *C. mimosae* | *C. nigriceps* | *T. penzigi* | None<sup>1</sup> | Other<sup>1</sup> |
| --- | --- | --- | --- | --- | --- | --- | --- |
| 1. Suyian Ranch | NE | 1.15 | 1.21 | 1.33 | 1.10 | 1.25 | 1.13 |
| 2. Mpala R.C. | NE | 2.30 | 2.51 | 2.07 | 1.84 | 3.10 |  |
| 3. Kitengela | SE |  | 1.20 | 1.28 | 1.42 | 1.17 | 1.87 |
| 4. Ngong Hills<sup>3</sup> | SE |  | 2.52 | 2.49 | 3.84 | 2.94 |  |
| 5. Isinya Road<sup>3</sup> | SE |  | 1.57 | 1.38 | 1.38 |  | 1.22 |
| 6. Mogotio | NW |  |  |  | 2.50 |  | 4.01 |
| 7. Koriema | NW |  |  | 5.28 | 4.64 | 3.00 |  |
| 9. Gilgil | SW |  |  | 1.50 | 1.58 | 1.53 |  |
| 10. Lake Naivasha | SW |  |  |  | 1.62 | 1.81 |  |
| 11. Narok Road | SW |  | 0.83 | 0.89 | 0.86 | 0.82 |  |

<sup>1</sup> Trees occupied by “none” indicate trees with no mature colonies or a few very tall trees for which we could not determine the canopy occupant. “Other” includes trees occupied solely by a *Lepisiota* species (n=29, all at Mogotio), as well as those trees with more than one species of ant in the canopy (n=21).
<sup>2</sup> Height and diameter measurements come from 69 trees in a 2400 m<sup>2</sup> subset of this area.
<sup>3</sup> At Ngong Hills and Isinya Road, we sampled from two separate transects within a short distance of each other. Both locations are given, but we pooled the height and ant occupancy data.
<sup>4</sup> At Mogotio, trees were occupied by a *Lepisiota* sp., or by *Lepisiota* and *T. penzigi* together, or in one case, by *Lepisiota*, *T. penzigi*, and *C. nigriceps*.
<sup>5</sup> At Lake Naivasha, only *T. penzigi* was found at the site we surveyed. However, we found *C. nigriceps* and a single colony of *C. mimosae* nearby along the South Lake Road.

**Figure 1:**
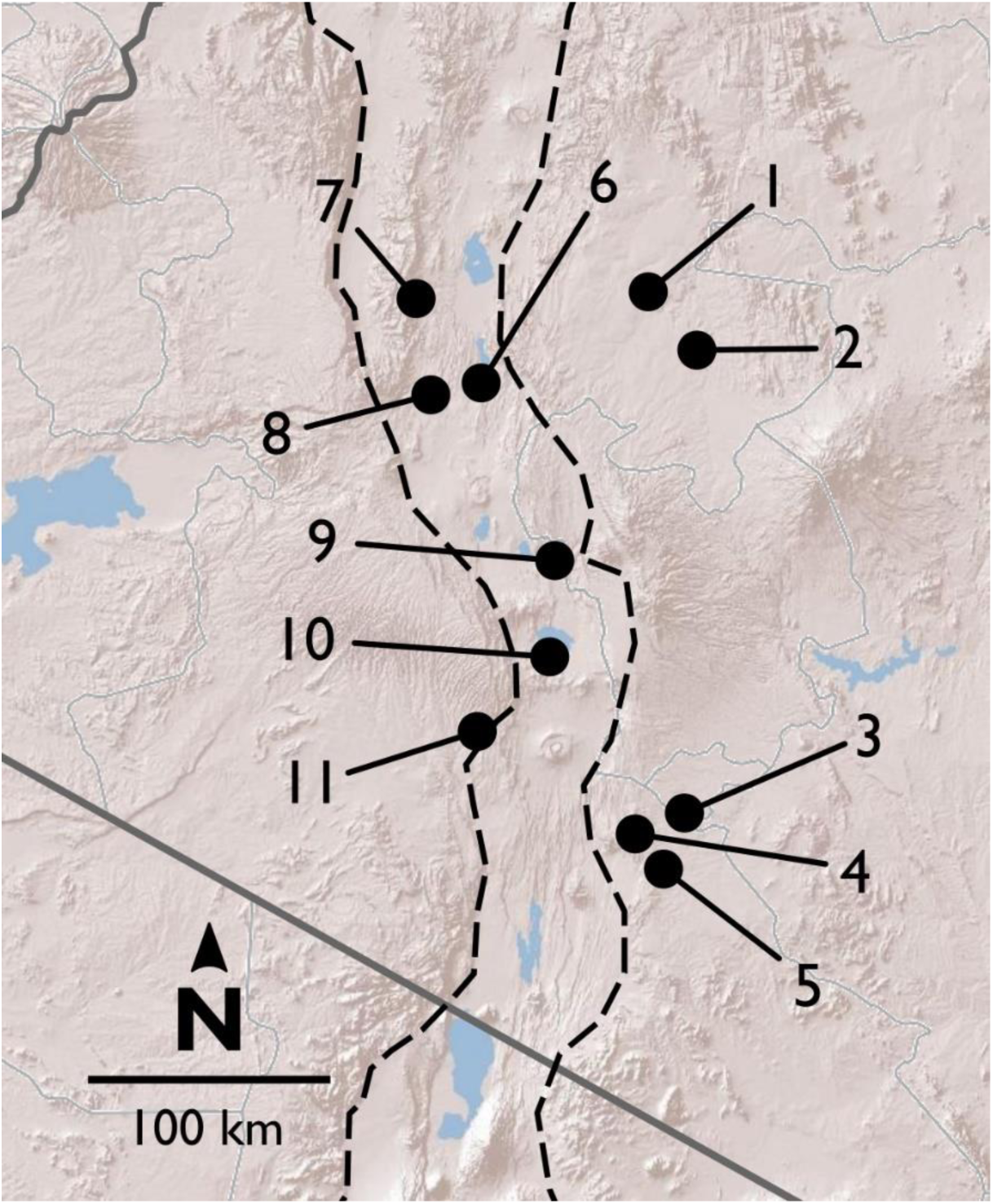
Map of field sites in and around the Rift Valley in central and southern Kenya (Rift shown by dashed red lines). Terrain image: Esri. *Northern highlands east of the Rift* 1. Suyian Ranch 2. Mpala Research Centre *Southern highlands east of the Rift* 3. Kitengela 4. Ngong Hills 5. Isinya Road *In and around the Rift: North* 6. Mogotio 7. Koriema 8. Mogotio-Koriema Road (population genetic samples only) *In and around the Rift: South* 9. Gilgil 10. Lake Naivasha 11. Narok Road (no population genetic samples)

At most of the sites, we surveyed a transect at least 0.15 ha in area, measuring every tree greater than 1.0 cm diameter at 0.5 meters above the ground. At Koriema and Gilgil, the patch of undisturbed habitat containing *V. drepanolobium* was smaller than this, so we surveyed every *V. drepanolobium* tree in these locations, and then calculated the area they occupied. Within these transects, we recorded the height, stem diameter at 0.5 meters above the ground, and ant occupant of each tree. At the Mpala site, we surveyed 6.7 hectares of the Mpala CTFS-ForestGeo plot for tree number and ant occupant, and a 0.24 ha subset of that area for tree number, ant occupant, and tree height and diameter. When the canopies of two trees were entangled, we counted this as a single tree for our purposes, since ants could move freely between them without having to leave the canopy; in these cases, and when a single tree had multiple stems 0.5 meters above the ground, we recorded the diameter of the single largest stem, and the height of the tallest point in the combined canopy.

### Collections

Between 2012-2014, mainly at the sites described above, we collected tissue from *V. drepanolobium* and its three primary ant associates, *C. mimosae*, *C. nigriceps*, and *T. penzigi*. These sites included five sites in the highlands east of the Rift Valley, and five in or adjacent to the Rift (we were not able to include a sixth Rift site, Narok, in the population genomic analysis because we surveyed this site after the other sites, in 2016, but we include the demographics for this site; for sample sizes see File S1). We did not include a fourth ant associate, *Crematogaster sjostedti*, in our population genetic study because we found it on *V. drepanolobium* trees only in Laikipia, at two adjacent sites, Suyian Ranch and Mpala Research Centre. *V. drepanolobium* leaf samples were dried and stored in desiccant, while workers of each species of ant were collected into 95-100% ethanol or into a buffered salt solution. *C. mimosae* and *C. nigriceps* colonies have been reported to spread across multiple trees (average colony size: 4.5 and 2.5 trees, respectively, at Mpala, Palmer *et al.* 2008). To avoid collecting multiple samples from a single colony spread across multiple trees, wherever possible, we collected workers of the same species from trees at least 5 meters apart. The fourth commonly studied ant associate, *C. sjostedti* Mayr, is found on *V. drepanolobium* only at sites in Laikipia (see Table 1), although it is free-living and facultative in its association with *V. drepanolobium*. It can be found associated with other tree species in areas such as the Tanzanian coast (Cochard *et al.* 2008). Since it is relatively rare on *V. drepanolobium* throughout Kenya, it was not included in the population genomic study.

### DNA extraction and sequencing

We extracted DNA from individuals (either a single worker for the ants, or a small number of leaflets for *V. drepanolobium*) using an AutoGenprep 965 Tissue/ES Cell DNA Extraction Kit (AutoGen). We used the Mouse Tail protocol for animal tissue. Throughout our protocols, we followed the manufacturer’s recommended protocols except where described. For extractions of *V. drepanolobium* gDNA, we modified this to lyse the cell walls by performing the lysis step using the CTAB buffer of Cullings (1992), to which was added 5 µL/mL β-mercaptoethanol. Genomic DNA was stored at −20° C before use.

The amount of genomic DNA was then increased by whole genome amplification, using Multiple Displacement Amplification via the REPLI-g mini kit (Qiagen) in 15 or 20 µL reactions.

We used the double-digest restriction-site associated DNA sequencing (RADseq) protocol of Peterson *et al*. (2012), but modified their protocol in the following ways: we started with an (amplified) genomic DNA mass of 150 ng, which we then digested with restriction enzymes chosen to provide hundreds to thousands of markers in each species. We used EcoRI-HF and BfaI (for the ants) or EcoRI-HF and MspI (for *V. drepanolobium*) (New England Biolabs). Bead cleanups throughout the protocol were performed with a MagNA bead solution which we made following the description of Rohland and Reich (2012). For bead cleanups, we added 1.5X volume of MagNA beads, but otherwise followed the protocol provided with Agencourt AMPure beads. We used the 48 inline indices for EcoRI described in the Sequences-S1 spreadsheet in the supplement of Peterson *et al*. (2012). We chose a range of 264-336 bp for the size selection step, which we performed using 2% ethidium bromide cassettes on a Sage Science Pippin Prep machine. The final PCR was set for 10 cycles.

These libraries were then sequenced in 100 bp, single-end reads on an Illumina HiSeq 2000 and 2500 at the Harvard University Bauer Core Facility.

### DNA sequence alignment and base-calling

To demultiplex the Illumina libraries, as well as to align reads across worker ants and call single nucleotide polymorphisms (SNPs), we used the program Stacks version 1.21 (Catchen *et al.* 2011, Catchen *et al.* 2013). For software analyses, we used the default parameters except where described. Reads were demultiplexed using the *process_radtags* function of Stacks, rescuing barcodes and RAD-tags, and disabling checking if the RAD site was intact.

We quality filtered reads using the FASTX-Toolkit version 0.0.13 (http://hannonlab.cshl.edu/fastx_toolkit/). For each read, the first seven basepairs, including the EcoRI-HF restriction site and two often-low-quality bases, were removed using the *fastx_trimmer* tool. The trimmed reads were then quality-filtered using the *fastq_quality_filter* tool, removing any reads with a quality score of less than 25 at more than 2% of bases.

We then aligned all reads for all individuals within each species using the *denovo_map.pl* script of Stacks, allowing 2 mismatches between loci when processing a single individual, and 2 mismatches when building the catalog. We also explored several other values for these parameters; the values given above gave the best combination of a larger number of alleles without large increases in heterozygosity which could indicate that different loci were being improperly combined (see File S2 for details). To build the final matrix of SNPs, we culled individuals for which sequencing had failed or had produced too low coverage to be useful, removing roughly a third of the individuals of each species. We called SNPs using the *populations* program of Stacks. We adjusted the stacks “-r” parameter to produce a similar number of markers (around 1000) within each species, but with variable amounts of missing data for each species. The -r parameter defines what proportion of the individuals within a species must have a given SNP for that SNP to be included in the data set. We filtered out sites with an unusually high heterozygosity (i.e., over 0.6), required a minimum minor allele frequency of 0.02, and outputted only a single, randomly-selected SNP per RADseq fragment. Although individuals varied in the degree to which they had missing data, individuals with high and low amounts of missing data were distributed fairly evenly across sites (File S3).

We also explored the effects of missing data on our results by producing two additional SNP data sets: In one, the “r” parameter was kept constant at 0.6 across all species, producing a variable number of SNPs for each species, but similar amounts of missing data. We used the same individuals as in the first data set. In the final data set, we used a subset of the individuals from the first two data sets, and included SNPs only if they were found in 100% of individuals within each species. This produced a matrix with no missing data. The results for these additional data sets were congruent with the original data set, and are presented in File S3.

### Population statistics

We divided the 10 sites into four populations based on their geography; these populations also broadly corresponded to the genetic clustering found by DAPC (described below). As described in Figure 1, these corresponded to northern and southern populations in and around the Rift, and northern and southern populations in the highlands east of the Rift. For *C. mimosae*, only a single colony was found in any of the Rift sites; we included this colony in the southern highlands population. We used Stacks to calculate observed and expected heterozygosity as well as θ (estimated as π). We calculated these values individually for each population, using the entire fragment for each locus identified above.

We used Arlequin 3.5.2.2 (Excoffier and Lischer 2010) to calculate summary statistics and pairwise F_ST_ between each population, after first translating Stacks’ genepop-format output files into Arlequin using PGDSpider 2.0.9.0 (Lischer and Excoffier 2012) in combination with a custom perl script. All such scripts used in this study are available on Dryad. Since we analyzed several data sets with varying degrees of missing data, we set Arlequin’s internal missing-data threshold to 1 (i.e., to include all loci with any data).

### Analysis of molecular variance (AMOVA)

We used an AMOVA to compare how much genetic variation was partitioned by the four geography-based populations (F_CT_) versus how much population-level variation was partitioned by the collection sites within each population (F_SC_). We used Arlequin to perform locus-by-locus AMOVAs, presenting the average F_SC_ and F_CT_ across all loci. We also report whether each measure is significantly greater than zero, as determined with 10,000 permutations.

### Genetic clustering analysis

To divide our individuals into genetic clusters, we used the adegenet 2.0.1 package (Jombart 2008, Jombart and Ahmed 2011), in R version 3.2.3 (R Core Team 2015). For each data set, we identified a number of clusters and assigned individuals to a single most likely cluster using find.clusters, with a maximum possible cluster number of 20, retaining all principle components, and selecting the cluster number with the lowest BIC score.

For *V. drepanolobium*, we tested whether trees assigned to a particular genetic cluster were more commonly occupied by colonies of a particular ant species. For each genetic cluster, we performed a multinomial exact test of goodness of fit test to determine whether the distribution of ant occupants within each genetic cluster was different from expected. To determine the expected ant-occupancy frequencies for each of the tree’s genetic clusters, we averaged the frequency of each ant species at the sites at which that cluster was found, weighting each site by the number of trees from that cluster that were present there. We considered only those sites where trees from multiple genetic clusters coexisted, and we tested association with four categories of ant occupant: *C. mimosae*, *C. nigriceps*, *T. penzigi*, and trees empty or occupied by a different species. When the expected value was zero for one of these species of ant occupant, we dropped that species from the test. We performed all tests using the xmulti function from the XNomial package in R (Engels 2015).

### Isolation by distance and environment

We then asked whether geographic structure was associated with environmental differences among sites, which might suggest local adaptation to abiotic conditions. To distinguish between isolation by environment and isolation by distance, we used the Multiple Matrix Regression with Randomization approach of Yang (2013), as implemented in the MRM function of the R package ecodist (Goslee and Urban 2007), testing significance with 1000 permutations. We characterized the environment using the bioclimatic variables for each site from the WorldClim database at 10 degree-minutes precision (Hijmans *et al*. 2005). Following He *et al.* (2016), we created clusters of climatic variables, such that each variable had a correlation of > 0.9 (or < −0.9) with at least one other variable in that cluster, and with no variables outside the cluster. We then chose one exemplar variable from each cluster, selecting Annual Mean Temperature, Temperature Annual Range, Annual Precipitation, Precipitation of the Warmest Quarter, Precipitation of the Wettest Quarter, and Precipitation of the Driest Quarter. In the case of the Lake Naivasha site, specimens were collected from five nearby locations along the South Lake Road, which differed in their WorldClim variables. We therefore treated them as separate sites for this analysis, despite their relative geographic proximity. For each species, the climate data from all sites were then scaled to have a mean of zero and standard deviation of one; environmental distance was then estimated as the Euclidian distance between each site for these six variables. Geographic distances between each site were calculated using the distGeo function from the geosphere package (Hijmans 2016). Genetic distances between each site were calculated using Nei’s distance, as implemented by the dist.genpop function of adegenet.

## Results

### Site survey

Plots varied substantially along several axes (Table 1, Figure 2). Tree density varied about five-fold, from 220 *V. drepanolobium* trees per hectare at Gilgil to a maximum of 1090 at the Isinya Road site; trees generally grew more densely in the eastern highlands than in the sites within the Rift. Average tree height also varied; at most sites the average height was 1-3 m, but at Koriema and Mogotio in the northwestern part of its range, *V. drepanolobium* trees averaged 4-5 meters, and grew up to 10 meters in height.

**Figure 2:**
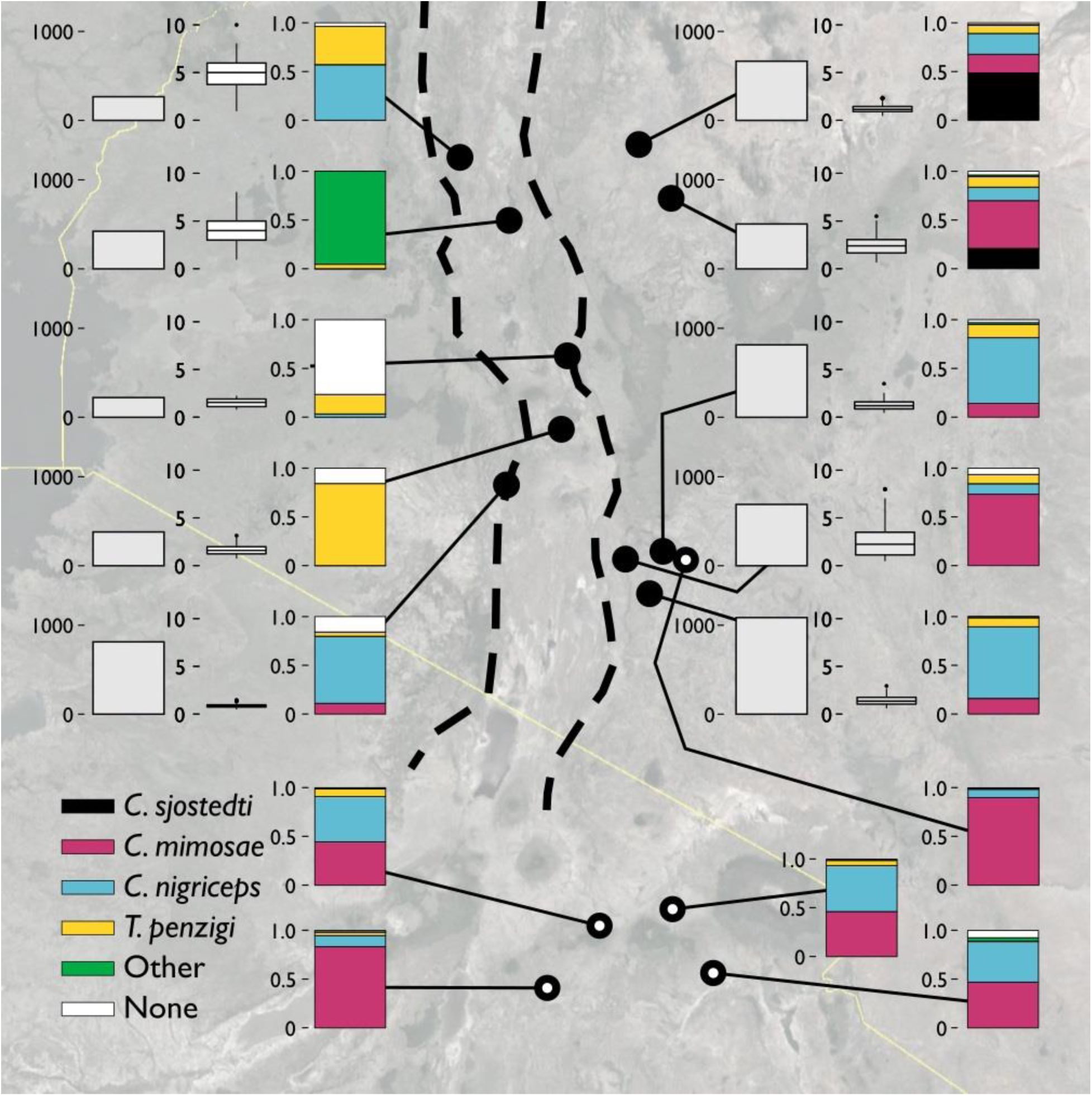
*The V. drepanolobium* ant-plant symbiosis varies across Kenya. Three plots summarize the characteristics of each site. The leftmost bar shows tree density, in trees/hectare. The middle box and whisker plot shows the distribution of tree heights (in meters) at each site. The rightmost bar shows the proportions of those trees occupied by each species of ant. “Other” indicated trees occupied by another species of ant, e.g., *Lepisiota* sp at Mogotio, or by more than one species of ant. “None” indicates that no colony appeared to occupy the tree. Sites marked by hollow circles are from Hocking (1970). Map image: Google, Landsay/Copernicus, US Dept. of State Geographer, Data SIO, NOAA, U.S.Navy, NGA, GEBCO.

Likewise, there was substantial variation in ant communities among sites. *C. sjostedti* was only found on *V. drepanolobium* trees in the northeastern sites of Suyian and Mpala. *C. mimosae* was more widespread, but commonly found only in the Eastern Highlands sites. In and around the Rift, *C. mimosae* was found rarely at Narok, and outside of that site, only a single colony was found, near the Lake Naivasha site. *C. nigriceps* and *T. penzigi*, however, were found at every location throughout the studied range. Furthermore, which ant was numerically dominant varied as well: for all four ant species, there was at least one site at which that species occupied more trees than did any other ant species.

As we have a single time point for each site, we are unable to measure directly interspecific dynamics such as the competitive hierarchy among the ants. However, we were able to analyze an indirect measurement, namely, the average height of trees occupied by each species. Previous work at Mpala has described how ant succession over the life span of the trees results in a characteristic pattern, where the average size of the trees occupied by each species follows the competitive hierarchy, with the most competitive *C. sjostedti* colonies are on the largest trees, and so on, down to the least competitive *T. penzigi* colonies, which are on the smallest trees (Young *et al.* 1997, Palmer *et al.* 2010).

We found a roughly similar pattern at our site in Mpala, with average tree heights following the competitive hierarchy (Figure 3), except that *C. mimosae* occupied larger trees than the more competitive *C. sjostedti*. In this case, however, the standard error of the mean for *C. sjostedti* was quite large, so the true population mean may easily have been greater than that of trees occupied *C. mimosae*. Moreover, competitive dynamics are known to vary even within Mpala (Palmer 2002), so this may be a result of our survey taking place in a different year and a different location within Mpala than the previous surveys.

**Figure 3:**
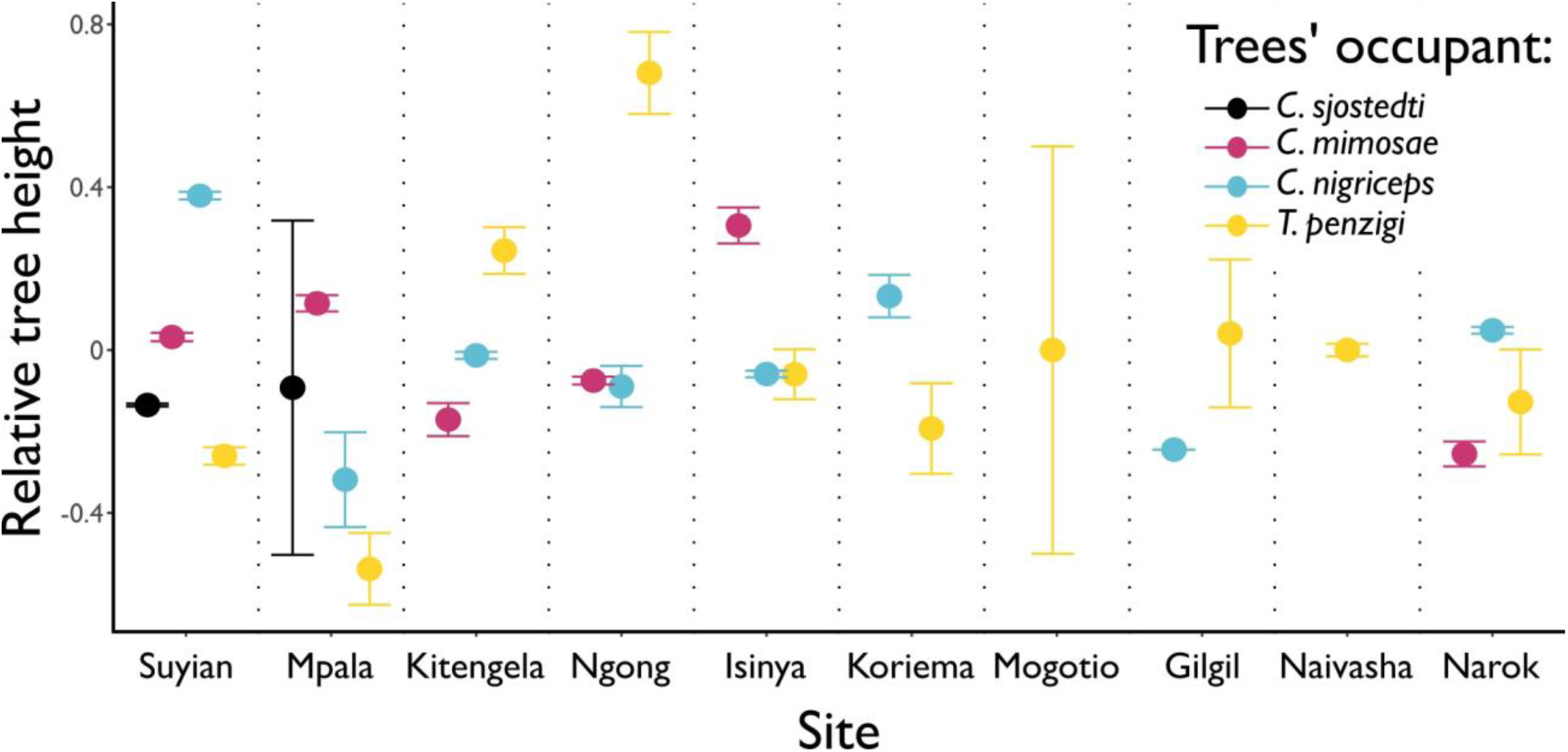
Different ants occupy the tallest trees at different sites. Points show the average height of the trees occupied by each ant species, at each height. Tree heights were standardized within sites to have a mean of 0 and standard deviation of 1. Whiskers show standard error. Each ant species occupies the tallest trees at one or more sites.

We did not find, however, that a similar pattern was maintained across all sites. At many sites, the largest trees were occupied by less competitively dominant ants: at the Ngong Hills, for instance, the largest trees were occupied by *T. penzigi*, while at Kitengela, the relationship between tree height and ant competitiveness was entirely reversed from the pattern at Mpala. These patterns in data may indicate that, just as the competitive hierarchy varies from site to site at Mpala, it also varies widely across the landscape. It may also be the case that the competitive hierarchy at the other sites follows the canonical hierarchy, but that what constitutes a valuable tree varies across sites. For example, *C. sjostedti* and *C. mimosae* occupy the most valuable trees, which are the largest trees at Mpala, but at other locations, the most valuable trees to the ants may be the medium-sized ones. Either case strongly suggests that the dynamics of interspecific competition among the ants is context dependent and varies strongly across the landscape.

### DNA sequence alignment and base-calling

We successfully genotyped individuals of each species at between 1027 and 1122 loci, depending on the species (Table 2).

**Table 2:**
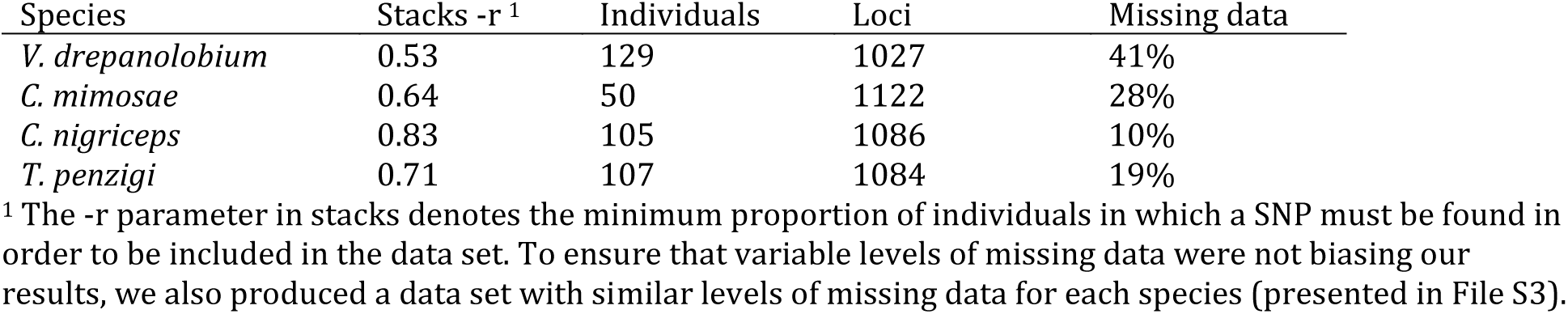
results of RADseq sequencing and base-calling

### Population statistics

*V. drepanolobium* populations generally have values of θ similar to those of the ants, suggesting similar effective population sizes across the four species studied here (Figure 4). θ values should be compared across species with caution, since θ is a product of effective population size multiplied by four times the mutation rate, which is unknown for the three ant species in question, or eight times the mutation rate for *V. drepanolobium*, which is tetraploid (Bukhari 1997). The estimates for F_ST_ for *V. drepanolobium* were relatively low, suggesting fewer barriers to gene flow for the tree than for its ant associates. Most of the structure in *V. drepanolobium* populations was between the Northern Rift (NW) population and the other three populations; F_ST_ values between the other three populations were much lower.

**Figure 4:**
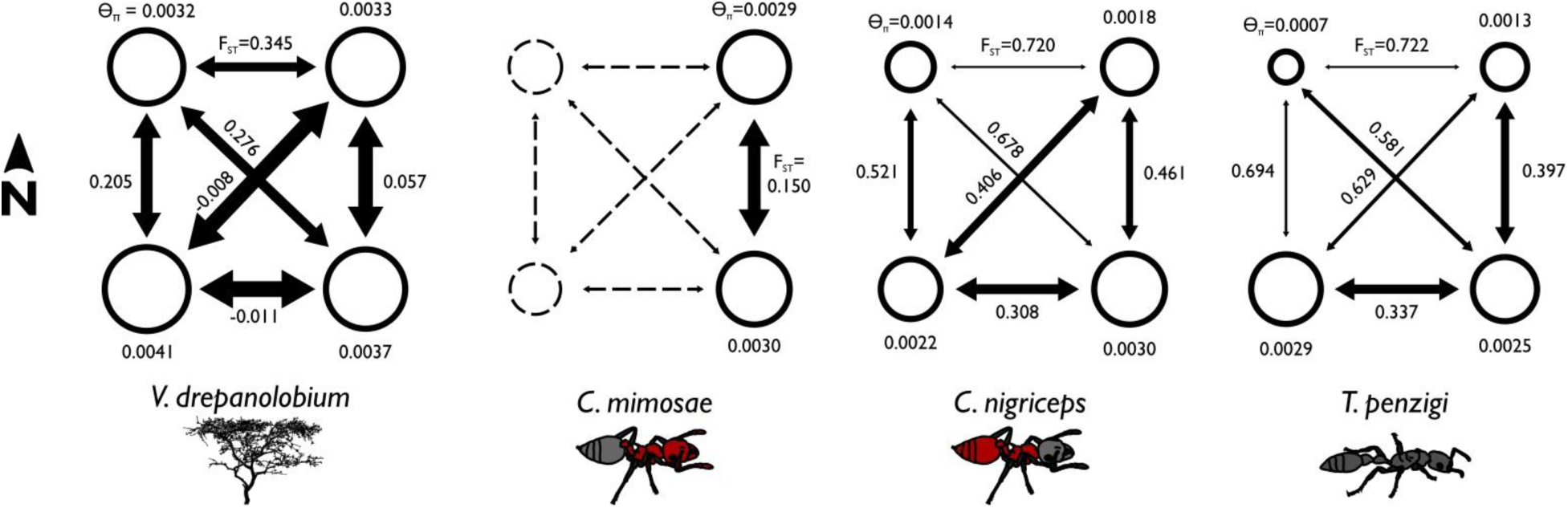
Summary statistics show differences in the population structure of the different species. The area of each circle is proportional to the θ_π_ value for each population, where θ_π_ is a measure of genetic diversity within populations that scales with effective population size. The arrows are scaled inversely to F_ST_, which measures the proportion of diversity unique to each population: lower F_ST_ values indicate greater gene flow between populations. θ values are similar across species, but F_ST_ values are lower for *V. drepanolobium* and *C. mimosae*, showing high gene flow. F_ST_ corresponds to the proportion of total genetic variation seen in the two populations that is partitioned between them. Values closer to zero indicate greater gene flow between the two populations. Negative values result from how F_ST_ values are averaged across multiple loci; they may be read as effectively zero.

Among the ants, these summary statistics suggested that *C. nigriceps* and *T. penzigi* have roughly similar population structures, and are both distinctly different from *C. mimosae*, which is the worst colonizer of the three at Mpala. *C. mimosae* spreads the least widely across the area studied, as it is only commonly found in the Eastern Highlands sites. However, it shows much less population structure between the two Eastern Highlands sites (F_ST_ = 0.15) than do the other two ants (F_ST_ = 0.461 and 0.396), suggesting that the Eastern Highlands region poses lower barriers to gene flow for *C. mimosae* than it does for *C. nigriceps* and *T. penzigi*.

The species also vary in terms of which populations contain the most diversity. In *V. drepanolobium* and *C. mimosae*, diversity values (θ_π_) are roughly similar among the different populations within each species; however, in *C. nigriceps* and *T. penzigi*, the southern populations have higher diversity values than the northern populations, and thus appear to be larger.

### Analysis of molecular variance

As shown in Table 3, AMOVA tests revealed significant genetic variation partitioned both among our geographic populations (F_CT_), and also among sites within those populations (F_SC_). This was true for all species. The species also varied in the magnitude of genetic variation partitioned among sites and populations. For *V. drepanolobium* and *C. mimosae*, both species have roughly similar F_SC_ and F_CT_ values. However, for *C. nigriceps* and *T. penzigi*, F_CT_ values were an order of magnitude higher than F_SC_ values. This suggests that a greater proportion of total genetic variation is partitioned by large-scale geography in the latter two species than in the former two species. In addition, the absolute value of F_SC_ was quite high for *T. penzigi*, suggesting that both inter-population variation and intra-population variation are important in this species.

**Table 3:**
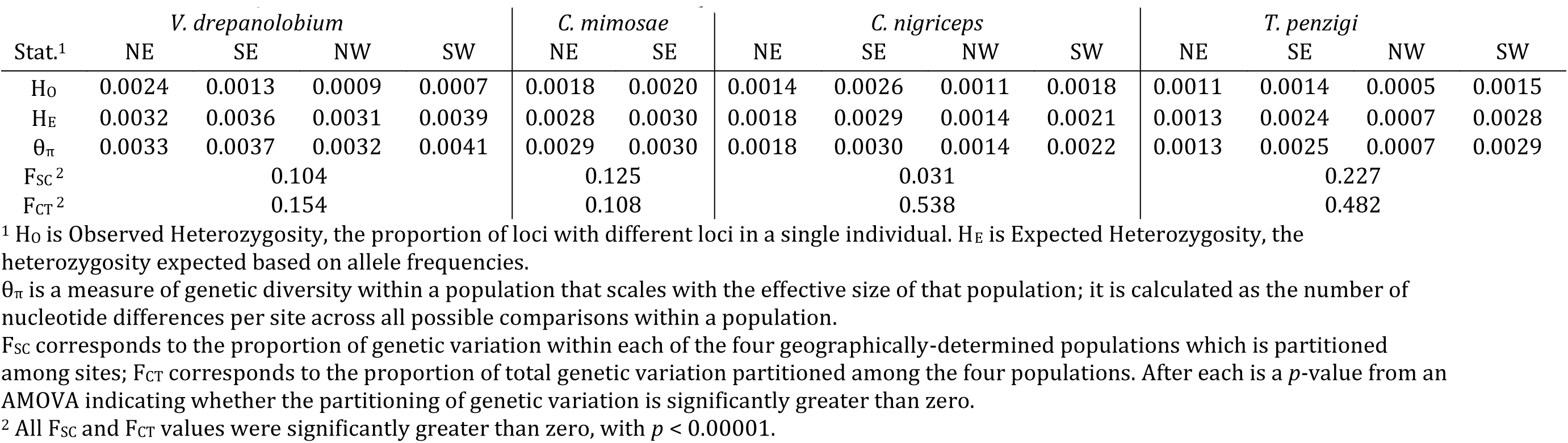
Population genetic summary statistics for each species.

**Table 4:** Pairwise FST between populations for each species

| Comparison | *V. drepanolobium* | *C. mimosae* | *C. nigriceps* | *T. penzigi* |
| --- | --- | --- | --- | --- |
| NE-SE | 0.057 | 0.150 | 0.461 | 0.397 |
| NE-SW | -0.008 |  | 0.406 | 0.629 |
| NE-NW | 0.345 |  | 0.720 | 0.722 |
| SE-SW | -0.011 |  | 0.308 | 0.337 |
| SE-NW | 0.276 |  | 0.678 | 0.581 |
| SW-NW | 0.205 |  | 0.521 | 0.694 |

### Genetic clustering analysis

For *V. drepanolobium*, the identified genetic clusters were weakly correlated with geography (Figure 5). Clustering shows separation between the sites in the Northern Rift and the sites in the Northern Highlands, with all the individuals in the Northern Highlands clustering together, and all of the individuals from the Northern Rift in a second cluster. However, in the Southern Highlands and Southern Rift sites, individuals belonging to both clusters could be found even at the same site.

**Figure 5:**
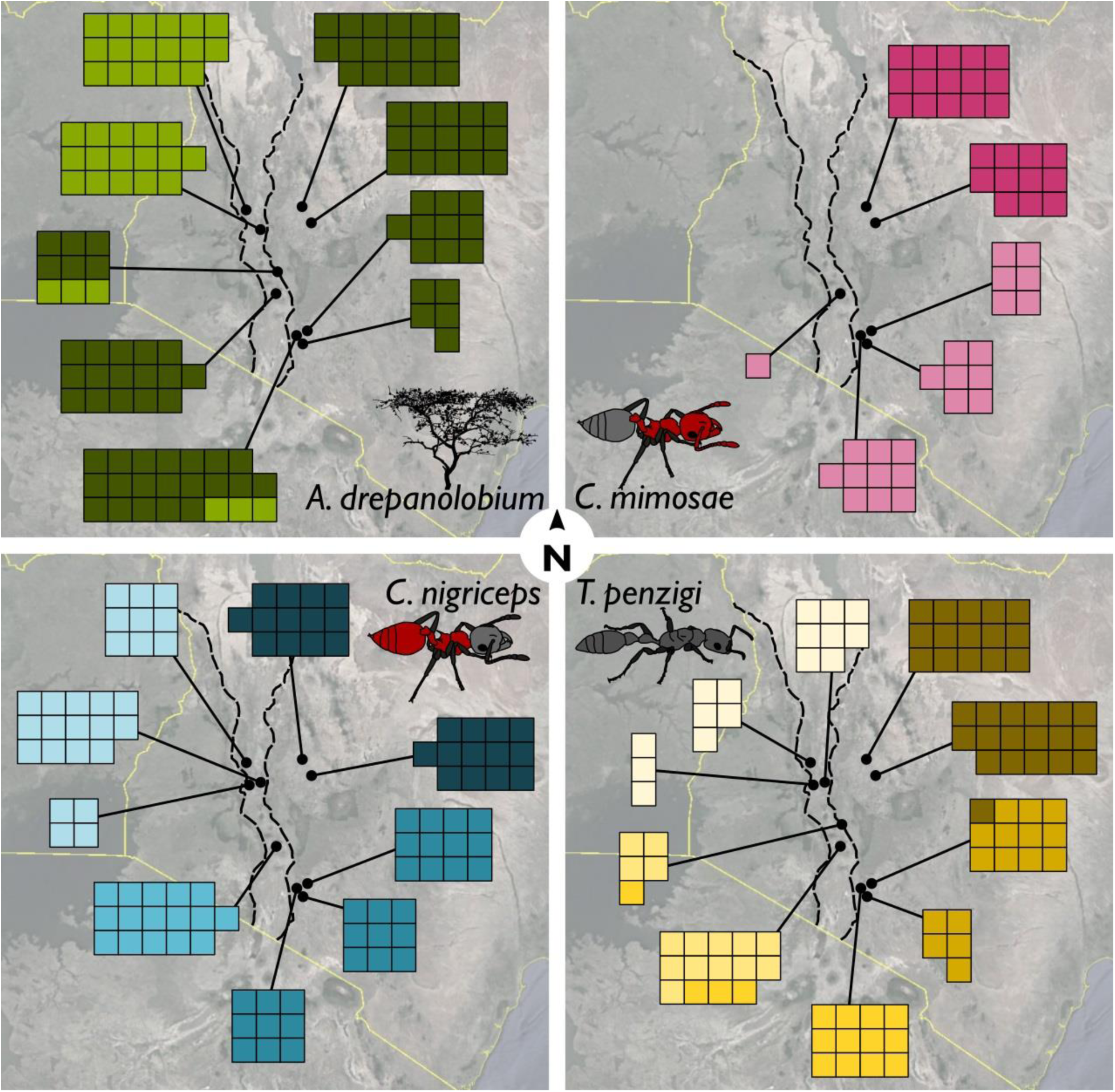
Ants showed more population structuring than did *V. drepanolobium*. Facets show the population clustering of (clockwise from upper left) *V. drepanolobium*, *C. mimosae*, *T. penzigi*, and *C. nigriceps*. Each individual was assigned to a single most probable cluster, which is indicated by the color of each individual (smaller squares). The total number of clusters was determined by taking the cluster number with the lowest BIC score. Map image: Google, Landsay/Copernicus, US Dept. of State Geographer, Data SIO, NOAA, U.S.Navy, NGA, GEBCO.

The ant species show a pattern that is quite different from that seen in their host tree: in these species, genetic clustering has a stronger association with geography in each case. Within each ant species, each of the four geographic areas (or two for *C. mimosae*) is dominated by a single cluster that is rarely, if ever, found outside that region. The exception to this is *T. penzigi*, where two genetic clusters are common in the Southern Rift sites, and the Ngong Hills site is dominated by one of these clusters, rather than clustering with the other Southern Highlands sites. Finally, the sole *C. mimosae* colony found in the Rift (at Lake Naivasha) clustered with the individuals from the Southern Highlands sites.

We found no evidence that trees assigned to different genetic clusters were associated with particular ant occupants (multinomial exact tests, *p* > 0.1 for all).

Population genetic statistics and clustering analyses indicate that the ant symbionts have particular population structures that differ both among themselves and between the ant symbionts and their host trees, indicating that each of the phytoecious ant species moves through the landscape in a way that differs from that of their host tree, and that the ant partners have different population histories as well. This finding was reinforced by a coalescent analysis, described in File S4.

### Isolation by distance and environment

For both *V. drepanolobium* and *C. mimosae*, we found no evidence of isolation by distance or isolation by environment (*p*-values > 0.05). In the case of *C. nigriceps*, we found evidence of both isolation by distance and isolation by environment (*p-*values < 0.05). For *T. penzigi*, we found evidence of isolation by distance (*p* < 0.05 in all three data sets), but no evidence of isolation by environment (*p* > 0.05 in all three data sets).

## Discussion

The *V. drepanolobium* system shows considerable variability across its range in central and southern Kenya. Much of this variation comes from comparing the sites in the Eastern Highlands to the sites in the Rift Valley, where the tree is less common, patches are smaller and sparser, and tree heights vary greatly. However, there is substantial variation even among the eastern highlands sites, where densities, average heights, and tree biomass vary by a factor of 2 or more. Nor is the variability confined to the plant; the different ant species show equally broad differences. In particular, the four ant species found at Mpala are commonly found together only in Laikipia; we did not find *C. sjostedti* outside of Laikipia, and we found *C. mimosae* only rarely outside of the Eastern Highlands (although it is also found at lower elevations in Tanzania: Hocking 1970). Even when comparing sites with the same species of ants, how the trees were divided among the ants varied: among our five highlands sites, numerical dominance varied, with the plurality, or even majority of trees being occupied by any one of *C. sjostedti*, *C. mimosae*, or *C. nigriceps*, depending on the site. We also found that the ant species occupying the tallest trees varied widely from site to site, suggesting that not just the proportions of species, but also, potentially, the outcomes of competition between them may vary from site to site. The variation we found across sites in Kenya agrees with what Hocking (1970) described for four sites in Tanzania and one in Kenya (Figure 2). He also found substantial difference across sites, with two of his sites dominated by *C. mimosae*, and three sites split about evenly between *C. mimosae* and *C. nigriceps*. His study and this study, combined, cover the bulk of the range in which *V. drepanolobium* is commonly found, and show that inter-site variation is the norm for this system.

The Rift Valley plays an important role in the genetics of other African acacias (Omondi *et al.* 2010, Ruiz Guajardo *et al.* 2010), and this is also true for *V. drepanolobium*: we observed low levels of gene flow between the Northern Rift population and the two highland populations for *V. drepanolobium*, *C. nigriceps*, and *T. penzigi.* However, the Rift as a barrier to gene flow does not explain the high F_ST_ between the two Rift sites in *V. drepanolobium*, or the relatively low F_ST_ between the Southern Rift and the highlands regions in *V. drepanolobium*, *C. nigriceps*, and *T. penzigi*. This suggests that the barrier effect of the Rift may vary across its length, or possibly that the Rift does not constitute a strong barrier to gene flow for this system, and some other feature of the landscape separates the Northern Rift population from the other three.

Population structure also varied widely among and between *V. drepanolobium* and its ant associates. Although all of the species showed substantial population structuring, *V. drepanolobium* had less structure than did its ant inhabitants: clusters of genetically similar individuals were spread across several different regions in Kenya, and most sites had individuals from more than one cluster, producing relatively low values of F_ST_. These clusters were not associated with particular ant inhabitants, and the overall pattern differed from the patterns observed for each of the different ant species, which exhibited much stronger geographic structuring. Most genetic clusters for the ant species were found only in a single region of Kenya, and comprised almost all of the individuals within that region, leading to relatively high F_ST_ values. The results of our coalescent analysis highlight the different population structures of each species, as the characteristics observed for each species were unlikely to be recovered if analyzed using the demographic parameters of any of the other species.

Variation in the population genetics of partners in a symbiosis is not unknown, especially in macrobe-microbe mutualisms (Werth and Sork 2010, Thornhill *et al.* 2013), but little evidence along these lines has been gathered for ant-plants or other macrosymbioses. In contrast to our results, Léotard *et al.* (2009) found congruent genetic signatures for the ant plant *Leonoardoxa africana* ssp. *africana* and its two ant symbionts, the defensive mutualist *Petalomyrmex phylax* and the non-defending *Cataulacus mckeyi*. Likewise, Blatrix *et al.* (2017) found largely congruent patterns between the ant plant *Barteria fistulosa* and its symbiont, *Tetraponera aethiops.* Blatrix *et al.* (2017) found that population structure was weaker in the ants than in the trees, whereas we found that population structure was stronger in the ants than in the trees. Clearly much work remains to determine whether any general patterns exist, but our results here suggest that the wide diversity of patterns seen in macrobe-microbe mutualisms (e.g., lichens) is also seen in large scale symbioses, like ant-plants.

We found little evidence that the genetic structure of *V. drepanolobium* and/or its associated ants is associated with environmental differences. Only for *C. nigriceps* did we find significant isolation-by-environment, suggesting that this ant species may be more locally adapted to environmental conditions than are the other species. It will be interesting to explore further to which environmental conditions particular populations of *C. nigriceps* are adapted. We also found little evidence of local coadaptation by partners of the symbiosis at each site. We discount this possibility for several reasons: first, the genetic clusters of the tree are not associated with particular ant species. Second, the genetic clusters of the tree are not distributed in the same pattern as the genetic clusters of the ants. This is perhaps not surprising, since individual trees (representing different lineages) of *V. drepanolobium* at Mpala have been well documented to interact with multiple different lineages of each ant species over the course of its lifetime (Palmer *et al.* 2010), reducing the opportunity for specialized coevolution between particular lineages of *V. drepanolobium* and its symbiotic ants. Furthermore, we find that different populations of *V. drepanolobium* include sites with extremely different ant composition. For instance, the sites at Suyian, Mpala, and Kitengela are dominated by a single genetic cluster of *V. drepanolobium*. However, each of these sites is dominated by a different ant species, and not all of the sites have the same ant species present. In light of this lack of local adaptation, it seems likely that all the partners in the symbiosis have experienced a wide array of different conditions at different places and times.

Many interspecific associations show variation across their geographic range and across different ecological conditions (Bronstein 1994, Thompson 2005), and the *V. drepanolobium* symbiosis appears to be an example of this kind of geographic mosaic. In light of the data presented here, it is clear that the dynamics of this association vary broadly across the Kenyan landscape. For instance, *T. penzigi* does not defend trees against large herbivores at Mpala (Palmer and Brody 2007) or Kitengela (Martins 2010); if this is true in the Rift as well, then *V. drepanolobium* at sites like Gilgil or Lake Naivasha, where *T. penzigi* dominates, are largely undefended against vertebrate herbivores. Likewise, if the competitive hierarchy is the same throughout the range, then *C. nigriceps* is a subordinate ant in the eastern part of the study area, but the dominant ant at sites in the Rift where *C. sjostedti* and *C. mimosae* are rare or absent. This discovery is the more striking because the *V. drepanolobium* system has been so well studied, producing many high-impact studies over the course of several decades (e.g., Stanton *et al.* 1999, Palmer *et al.* 2008, Palmer *et al.* 2010). If such a system can contain cryptic ecological variation across its range, then this is likely the case in other, as well-studied, model systems. This is particularly important for evolutionary studies: if we are to calculate the costs and benefits to an organism of particular traits, we must understand not only the current, local environment, but also the environments experienced by that organism’s lineage. Systems like *V. drepanolobium*, where single panmictic populations experience a wide variety of local conditions, force us to study the system across a number of different sites if we are to understand fully its ecological and evolutionary dynamics.

## Author Contributions

JHB and NEP conceived the experiments. JHB, DJM, and PMM collected specimens, site data, and provided logistical support in the field. JHB performed the DNA sequencing and analyses. JHB and NEP wrote the paper with the input of DJM and PMM.

## Acknowledgements

We thank the many people who helped with fieldwork, especially Ezekiel Chebii, Richard Childers, Stephen Gitau, Francis Irinye, Isaack Leting, Thomas Mojong, Julianne Pelaez, and Lori Shapiro. Many people showed us hospitality in Kenya, but particularly Anne Powys and Gilfrid Powys at the Suyian Ranch, Nani Croze and Eric Krystall at Kitengela, and the staff at the Mpala Research Centre. The RADseq technique, and much bioinformatic advice, were generously provided by Brant Peterson, Jesse Weber, and Hopi Hoekstra. Advice and support in the laboratory came from Ian Butler, Rachel Hawkins, Julianne Pelaez, Lukas Rieppel, and Maggie Starvish in the Pierce lab; and from Christian Daly, Claire Reardon, and Jennifer Couget at the Harvard Bauer Core Facility. Most of the computations in this paper were run on the Odyssey cluster supported by the FAS Division of Science, Research Computing Group at Harvard University. We are also thankful for advice on bioinformatics and analyses from Bruno de Medeiros, Seth Donoughe, Sarah Kocher, David Haig, Brian Farrell, and Scott Edwards; and to Noah Whiteman for his comments on the manuscript. Our research was supported by grants from the Putnam Expeditionary Fund of the Museum of Comparative Zoology, the Mind, Brain, Behavior Interfaculty Initiative at Harvard, FQEB grant no. RFP-12-06 from the National Philanthropic Trust and the National Science Foundation (NSF SES-0750480) to NEP. We dedicate this paper to the memory of our friend, Gilfrid Powys, who generously made his land available to us and supported our work in numerous ways throughout the duration of this research. He will be greatly missed.

## Data Accessibility

We will upload the DNA sequences used in this project to the NCBI SRA. Scripts used in these analyses will be available on Dryad.

## Supporting Material

**File S1: Sample sizes and climate data for each site in the population genetic survey**

**Table S1.1:** Sample sizes for population genomic analyses

| Site | Samples genotyped from each site for: |  |  |  |
| --- | --- | --- | --- | --- |
|  | *V. drepanolobium* | *C. mimosae* | *C. nigriceps* | *T. penzigi* |
| Suyian Ranch | 18 (12) | 15 (5) | 13 (5) | 15 (12) |
| Mpala R.C. | 15 (8) | 11 (4) | 13 (3) | 16 (12) |
| Kitengela | 10 (5) | 6 (4) | 12 (7) | 12 (6) |
| Ngong Hills<sup>1</sup> | 23 (5) | 10 (8) | 9 (8) | 12 (6) |
| Isinya Road<sup>1</sup> | 5 (0) | 7 (3) | 9 (7) | 5 (4) |
| Koriema | 18 (2) | 0 (0) | 9 (7) | 5 (3) |
| Mogotio | 16 (5) | 0 (0) | 14 (12) | 8 (7) |
| Mogotio-Koriema Road<sup>1</sup> | 0 (0) | 0 (0) | 4 (3) | 3 (0) |
| Gilgil | 9 (0) | 0 (0) | 0 (0) | 5 (3) |
| Lake Naivasha<sup>2</sup> | 10 (0) | 0 (0) | 0 (0) | 14 (4) |
| South Lake Road 1<sup>2</sup> | 0 (0) | 0 (0) | 6 (6) | 5 (1) |
| South Lake Road 2<sup>2</sup> | 0 (0) | 0 (0) | 4 (4) | 5 (4) |
| South Lake Road 3<sup>2</sup> | 2 (0) | 0 (0) | 6 (6) | 0 (0) |
| South Lake Road 4<sup>2</sup> | 4 (0) | 1 (1) | 6 (5) | 2 (0) |

**Table S1.2:**
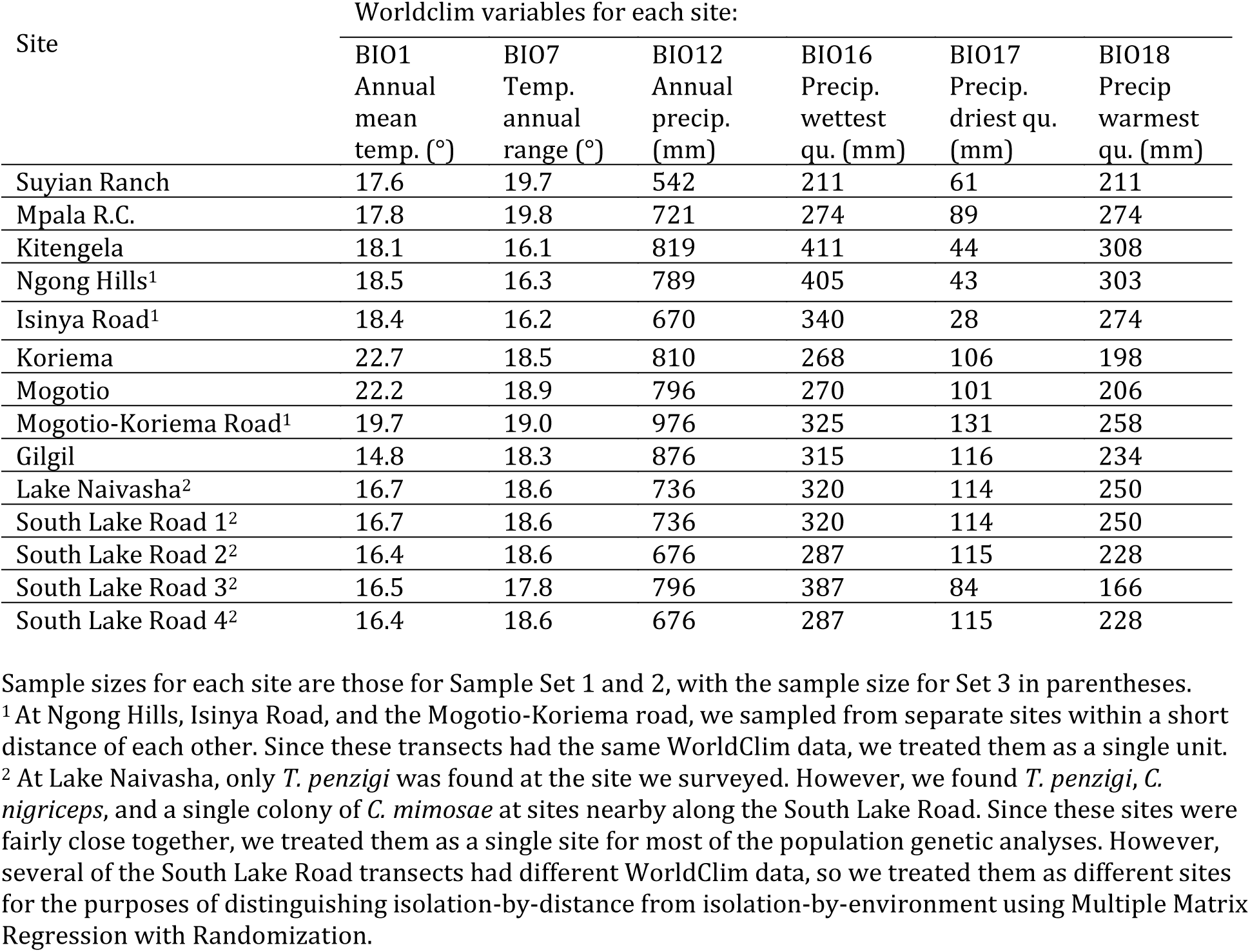
Worldclim data from population genomic sites

**File S2: Choosing stacks parameters**

When building the Stacks catalogs of loci, we allowed two parameters to vary: the number of mismatches allowed between loci when processing an individual (-M parameter), and the number of mismatches allowed between loci when building the catalog (-n parameter). To present the outcomes, we use the individuals and run parameters of Data Set 1 to call SNPs. We then calculated average heterozygosity of the SNPs using the adegenet package version 2.0.1 (Jombart 2008, Jombart and Ahmed 2011) in R version 3.2.3 (R Core Team 2015).

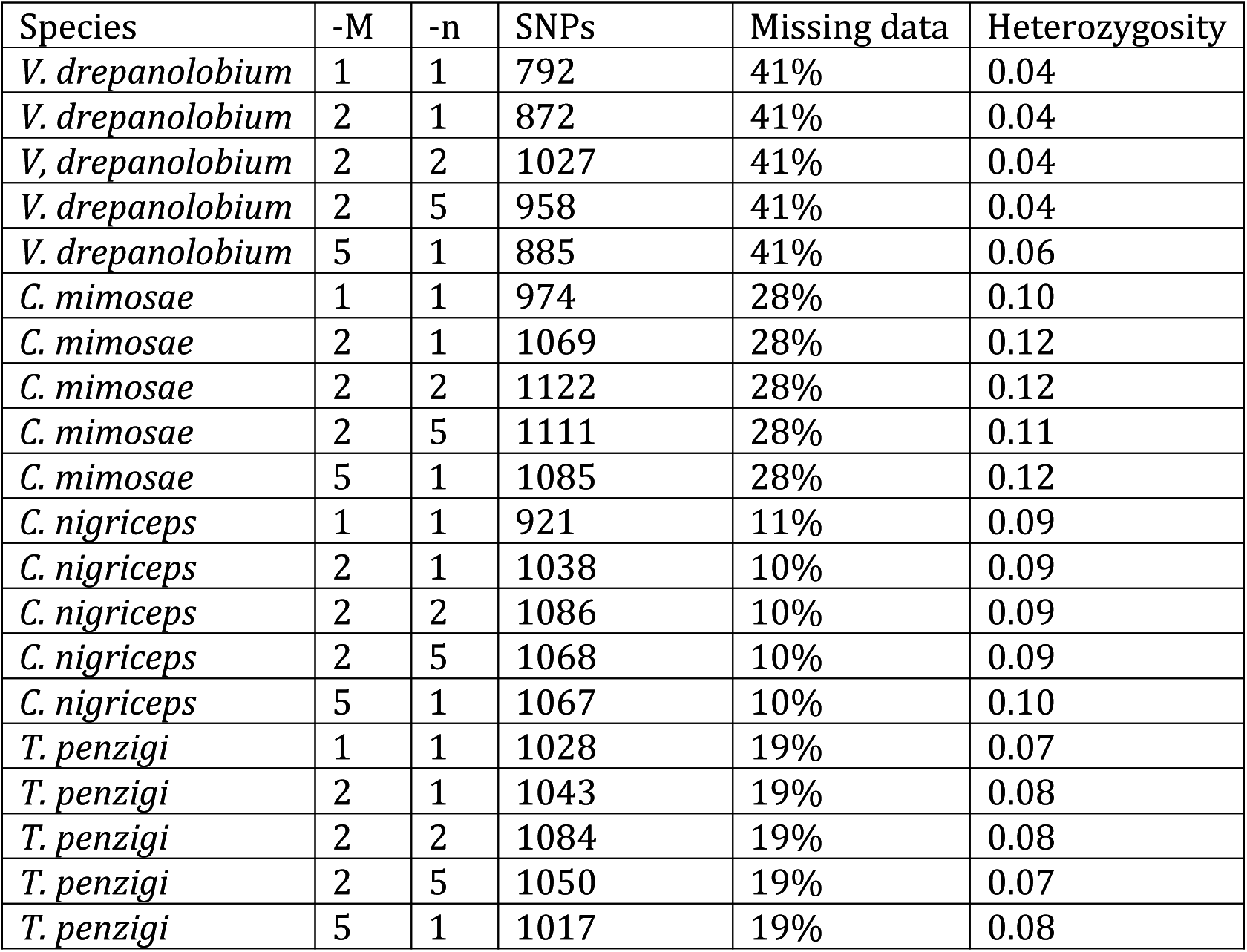

---

**File S3: Our results are not biased by missing data**

*Distribution of missing data*

Individuals varied widely in the amount missing data: most individuals had low levels of missing data, while a few individuals had considerably larger amounts (Figure S3.1). Fortunately, the individuals with larger amounts of missing data were quite evenly spread across populations; i.e., our results were not biased because some populations had disproportionate numbers of individuals with high missing data. To further test whether our results were affected by missing data, we also analyzed three data sets in parallel, each of which had different degrees of missing data (including no missing data in the final data set).

*DNA sequence alignment and base-calling*

As seen in Table S3.1, we successfully genotyped individuals of each species at between 120 and 3763 loci, depending on the species and the data set.

**Table S3.1:** results of RADseq sequencing and base-calling

| Species | Data set | Stacks -r <sup>1</sup> | Individuals | Loci | Missing data |
| --- | --- | --- | --- | --- | --- |
| *V. drepanolobium* | 1 | 0.53 | 129 | 1027 | 41% |
|  | 2 | 0.6 | 129 | 400 | 35% |
|  | 3 | 1 | 37 | 120 | 0% |
| *C. mimosae* | 1 | 0.64 | 50 | 1122 | 28% |
|  | 2 | 0.6 | 50 | 1386 | 30% |
|  | 3 | 1 | 25 | 613 | 0% |
| *C. nigriceps* | 1 | 0.83 | 105 | 1086 | 10% |
|  | 2 | 0.6 | 105 | 3763 | 23% |
|  | 3 | 1 | 73 | 463 | 0% |
| *T. penzigi* | 1 | 0.71 | 107 | 1084 | 19% |
|  | 2 | 0.6 | 107 | 1655 | 24% |
|  | 3 | 1 | 62 | 231 | 0% |

<sup>1</sup> The -r parameter in stacks denotes the minimum proportion of individuals in which a SNP must be found in order to be included in the data set.

*Population statistics*

The three data sets produced broadly similar results, despite differences in the individuals and SNPs making up each set (Tables S4.2, S4.3).

*Analysis of molecular variance*

As shown in Table S3.2, AMOVA tests revealed significant genetic variation partitioned both among our geographic populations (F_CT_), and also among sites within those populations (F_SC_). This was true for all species and all data sets.

**Figure S3.1:**
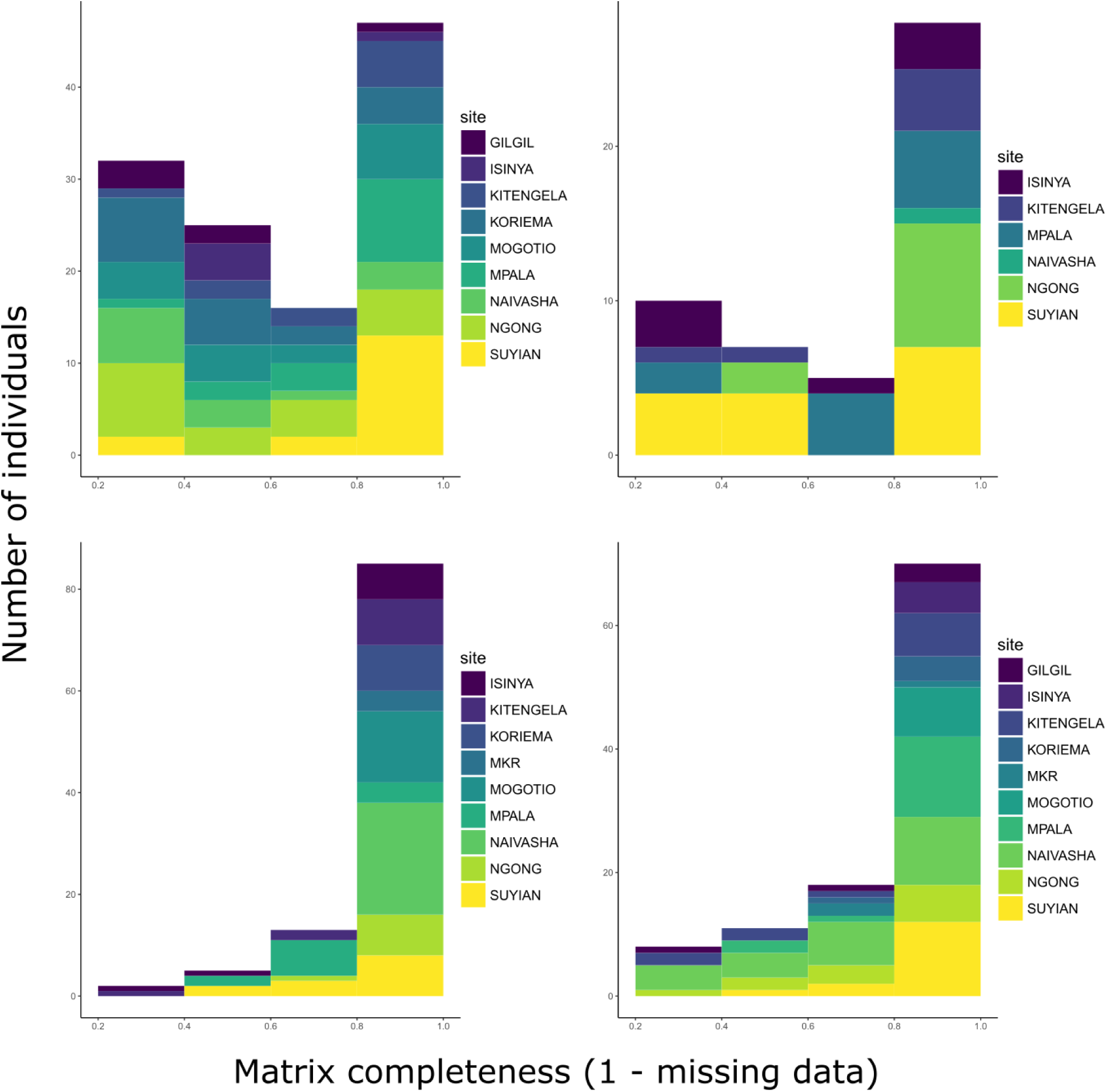
Levels of missing data vary across individuals. Histogram bars are colored according to the number of individuals in that class from each site. MKR stands for Mogotio-Koriema Road. Although individuals vary in their degree of missing data, individuals with large amounts of missing data are mostly evenly distributed across sites.

**Table S3.2:**
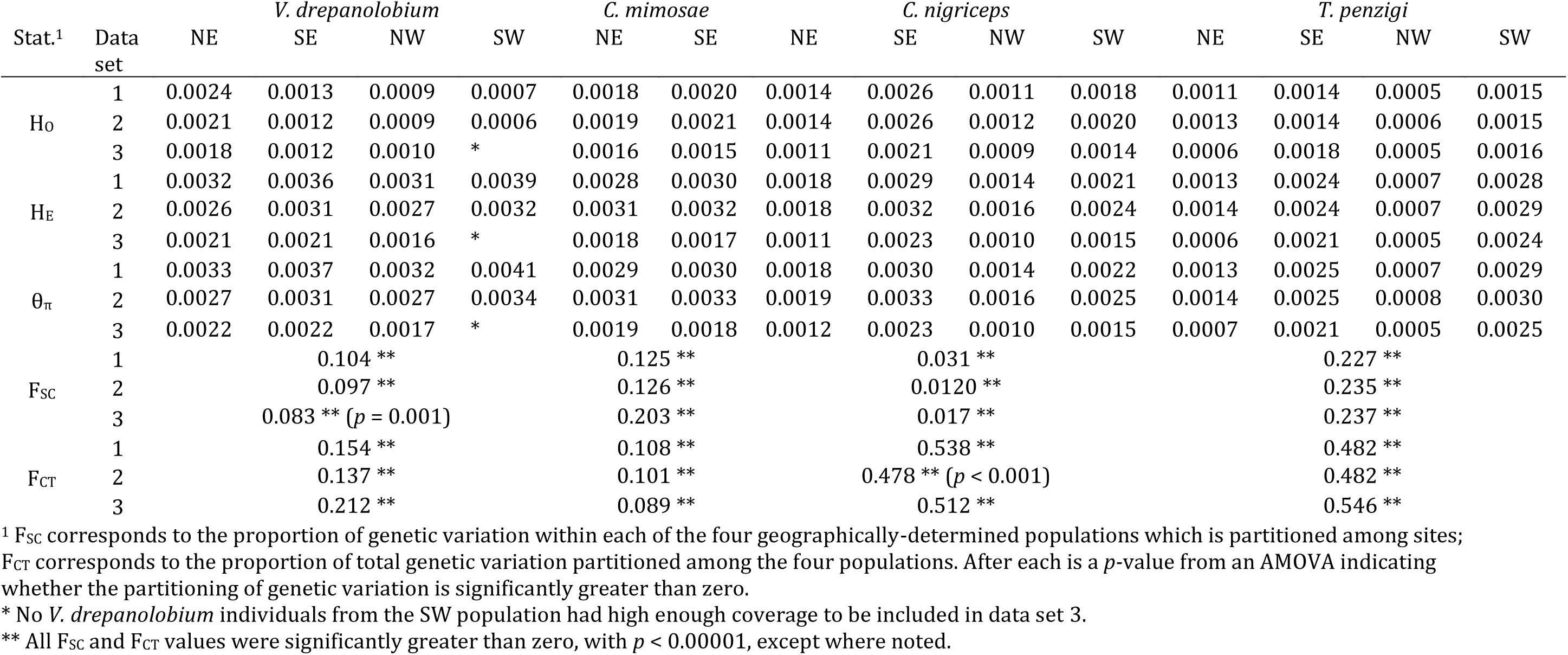
Population genetic summary statistics for each species.

| Stat.<sup>1</sup> | Data set | *V. drepanolobium* |  |  |  | *C. mimosae* |  | NE | *C. nigriceps* |  |  | NE | *T. penzigi* |  |  |
| --- | --- | --- | --- | --- | --- | --- | --- | --- | --- | --- | --- | --- | --- | --- | --- |
|  |  | NE | SE | NW | SW | NE | SE |  | SE | NW | SW |  | SE | NW | SW |
| H<sub>O</sub> | 1 | 0.0024 | 0.0013 | 0.0009 | 0.0007 | 0.0018 | 0.0020 | 0.0014 | 0.0026 | 0.0011 | 0.0018 | 0.0011 | 0.0014 | 0.0005 | 0.0015 |
|  | 2 | 0.0021 | 0.0012 | 0.0009 | 0.0006 | 0.0019 | 0.0021 | 0.0014 | 0.0026 | 0.0012 | 0.0020 | 0.0013 | 0.0014 | 0.0006 | 0.0015 |
|  | 3 | 0.0018 | 0.0012 | 0.0010 | * | 0.0016 | 0.0015 | 0.0011 | 0.0021 | 0.0009 | 0.0014 | 0.0006 | 0.0018 | 0.0005 | 0.0016 |
| H<sub>E</sub> | 1 | 0.0032 | 0.0036 | 0.0031 | 0.0039 | 0.0028 | 0.0030 | 0.0018 | 0.0029 | 0.0014 | 0.0021 | 0.0013 | 0.0024 | 0.0007 | 0.0028 |
|  | 2 | 0.0026 | 0.0031 | 0.0027 | 0.0032 | 0.0031 | 0.0032 | 0.0018 | 0.0032 | 0.0016 | 0.0024 | 0.0014 | 0.0024 | 0.0007 | 0.0029 |
|  | 3 | 0.0021 | 0.0021 | 0.0016 | * | 0.0018 | 0.0017 | 0.0011 | 0.0023 | 0.0010 | 0.0015 | 0.0006 | 0.0021 | 0.0005 | 0.0024 |
| θ<sub>π</sub> | 1 | 0.0033 | 0.0037 | 0.0032 | 0.0041 | 0.0029 | 0.0030 | 0.0018 | 0.0030 | 0.0014 | 0.0022 | 0.0013 | 0.0025 | 0.0007 | 0.0029 |
|  | 2 | 0.0027 | 0.0031 | 0.0027 | 0.0034 | 0.0031 | 0.0033 | 0.0019 | 0.0033 | 0.0016 | 0.0025 | 0.0014 | 0.0025 | 0.0008 | 0.0030 |
|  | 3 | 0.0022 | 0.0022 | 0.0017 | * | 0.0019 | 0.0018 | 0.0012 | 0.0023 | 0.0010 | 0.0015 | 0.0007 | 0.0021 | 0.0005 | 0.0025 |
| F<sub>SC</sub> | 1 |  | 0.104 ** |  |  | 0.125 ** |  |  | 0.031 ** |  |  |  | 0.227 ** |  |  |
|  | 2 |  | 0.097 ** |  |  | 0.126 ** |  |  | 0.0120 ** |  |  |  | 0.235 ** |  |  |
|  | 3 |  | 0.083 ** (*p* = 0.001) |  |  | 0.203 ** |  |  | 0.017 ** |  |  |  | 0.237 ** |  |  |
| F<sub>CT</sub> | 1 |  | 0.154 ** |  |  | 0.108 ** |  |  | 0.538 ** |  |  |  | 0.482 ** |  |  |
|  | 2 |  | 0.137 ** |  |  | 0.101 ** |  |  | 0.478 ** (*p* < 0.001) |  |  |  | 0.482 ** |  |  |
|  | 3 |  | 0.212 ** |  |  | 0.089 ** |  |  | 0.512 ** |  |  |  | 0.546 ** |  |  |

<sup>1</sup> F<sub>SC</sub> corresponds to the proportion of genetic variation within each of the four geographically-determined populations which is partitioned among sites;
F<sub>CT</sub> corresponds to the proportion of total genetic variation partitioned among the four populations. After each is a *p*-value from an AMOVA indicating whether the partitioning of genetic variation is significantly greater than zero.
\* No *V. drepanolobium* individuals from the SW population had high enough coverage to be included in data set 3.
\*\* All F<sub>SC</sub> and F<sub>CT</sub> values were significantly greater than zero, with *p* < 0.00001, except where noted.

**Table S3.3:** Pairwise FST between populations for each species

| Comparison | Data Set | *V. drepanolobium* | *C. mimosae* | *C. nigriceps* | *T. penzigi* |
| --- | --- | --- | --- | --- | --- |
| NE-SE | 1 | 0.057 | 0.150 | 0.461 | 0.397 |
|  | 2 | 0.038 | 0.139 | 0.354 | 0.400 |
|  | 3 | 0.063 | 0.156 | 0.467 | 0.398 |
| NE-SW | 1 | -0.008 |  | 0.406 | 0.629 |
|  | 2 | -0.022 |  | 0.197 | 0.638 |
|  | 3 | * |  | 0.417 | 0.684 |
| NE-NW | 1 | 0.345 |  | 0.720 | 0.722 |
|  | 2 | 0.321 |  | 0.651 | 0.711 |
|  | 3 | 0.320 |  | 0.743 | 0.819 |
| SE-SW | 1 | -0.011 |  | 0.308 | 0.337 |
|  | 2 | -0.021 |  | 0.284 | 0.347 |
|  | 3 | * |  | 0.280 | 0.325 |
| SE-NW | 1 | 0.276 |  | 0.678 | 0.581 |
|  | 2 | 0.251 |  | 0.658 | 0.583 |
|  | 3 | 0.387 |  | 0.660 | 0.616 |
| SW-NW | 1 | 0.205 |  | 0.521 | 0.694 |
|  | 2 | 0.182 |  | 0.498 | 0.703 |
|  | 3 | * |  | 0.493 | 0.731 |

\* None of the *V. drepanolobium* individuals from the SW population had high enough coverage to be included in data set 3, and thus there are no pairwise comparison between the SW population and the others.

*Genetic clustering analysis*

For *V. drepanolobium*, the identified genetic clusters were weakly correlated with geography. All three data sets show separation between the sites in the Northern Rift and the sites in the Northern Highlands, with all the individuals in the Northern Highlands clustering together, and all of the individuals from the Northern Rift in a second cluster. However, in the Southern Highlands and Southern Rift sites, individuals belonging to both clusters (as well as a third cluster in data set 2) may be found even at the same site.

Each of the ant species shows a much stronger associations between genetic clustering and geography. Across all data sets, each of the four geographic areas was dominated by a single cluster that was rarely, if ever, found outside that region. The exceptions to this were *C. nigriceps*, in which data set 2 showed two genetic clusters in the Northern Highlands, and *T. penzigi*, where two genetic clusters were common in the Southern Rift sites, and the Ngong Hills site was dominated by one of these clusters, rather than clustering with the other Southern Highlands sites. In data set 3, the Southern Rift sites and the Ngong Hills formed a single cluster, and the other two Southern Highlands sites were each dominated by their own cluster. Finally, the representative individual from the sole *C. mimosae* colony found in the Rift (at Lake Naivasha) clustered with the individuals from the Southern Highlands sites.

We found no evidence that trees assigned to different genetic clusters were associated with particular ant occupants (multinomial exact tests, *p* > 0.1 for all); this equally the case for all three data sets.

*Isolation by distance and environment*

Within each of the four species, the three data sets produced similar results. For both *V. drepanolobium* and *C. mimosae*, we found no evidence of isolation by distance or isolation by environment in any of the data sets (all *p*-values > 0.05). In the case of *C. nigriceps*, we found evidence of both isolation by distance and isolation by environment (all *p-*values < 0.05), except for data set 3, in which we only found evidence for isolation by distance. For *T. penzigi*, we found evidence of isolation by distance (*p* < 0.05 in all three data sets), but no evidence of isolation by environment (*p* > 0.05 in all three data sets).

**File S4: Coalescent analyses provide further evidence of distinct population structures for each species**

*Coalescent analysis*

To quantify the degree to which the four mutualists moved through the landscape in different ways, we used fastsimcoal2.5.2.21 (Excoffier and Foll 2011, Excoffier *et al.* 2013), which uses a coalescent approach to estimate population parameters from site frequency spectra (SFS), which summarize the distribution of minor allele frequencies within and between populations. We used Arlequin to produce site frequency spectra for each species, using data set 3, since this requires a data matrix without missing data. The stacks data set used to produce site frequency spectra includes only single-nucleotide polymorphisms, and not sites that are invariant across all populations. We calculated the number of these sites for each species as *L*_*v*_ * (89-(*S*/(*L*_*v*_ + *L*_*i*_)) / (*S*/(*L*_*v*_ + *L*_*i*_)), where *L*_*v*_ and *L*_*i*_ are the number of loci with and without any variable sites, respectively; *S* is the number of variable sites across all loci; and 89 is the length of the sequenced fragment. This value, rounded to the nearest integer, was added to the number of completely invariant sites for each site frequency spectrum.

These SFS were then used to estimate a number of parameters for each species, including effective population size, growth rates for each population, and migration rates between each population (see Figure S4.1). We used pairwise FST values for each species to determine which populations coalesced (for *V. drepanolobium*, the other populations coalesced with the Northern Highlands population; for *C. nigriceps*, with the Southern Highlands site; for *T. penzigi*, with the Southern Rift site). The following settings were used to estimate parameters for each run: 100,000 simulations were performed to estimate the expected derived SFS; a minimum and maximum of 10 and 20 conditional maximization cycles, respectively, were used to estimate parameters, with a stop criterion of 0.001. Initial parameters were drawn from a log-uniform prior from 10 to 10^9^ for population sizes, a uniform prior from 10 to 10,000 for divergence times, and a log-uniform prior from 10^-8^ to 10^-3^ for migration rates. We performed 100 runs for each species. We present here the results from all 100 runs, weighted by their likelihood using the Hmisc package in R (Harrell 2016).

Finally, we repeated the above analysis, but instead of drawing all population parameters from a prior distribution, we instead set population sizes and growth rates using the maximum-likelihood parameters from each species. We then compared the estimated likelihoods of these scenarios to show whether genes flowed through the landscape in similar ways for each species. For instance, we used fastsimcoal2 to estimate the likelihood of the *C. nigriceps* data given the most likely parameters for that species. We then estimated the likelihood of the *C. nigriceps* data, using instead the most likely parameters for *T. penzigi*. If the likelihoods in each case are similar, this suggests that the population structures of *C. nigriceps* and *T. penzigi* are similar; while if the *C. nigriceps* data is much more likely under the *C. nigriceps* parameters than under the *T. penzigi* parameters, this suggests that *C. nigriceps* and *T. penzigi* have distinctly different population structures.

The details for estimating these likelihoods are as follows: For each species, we calculated the ratio between each population size and between each individual migration rate under the most likely scenario produced above. This provided a characteristic population structure for each of the four species. We then repeated the previous runs, using the same parameters and priors, except that we estimated the likelihood of the data for each species under the characteristic population structure of each other species. To provide a comparison, we also estimated the likelihood of the data for a single species under its own characteristic population structure. When two species had different numbers of populations (i.e., *C. mimosae* has only two populations, while *C. nigriceps* has four), we reduced the data set of the species with more populations to allow the appropriate comparison.

**Figure S4.1:**
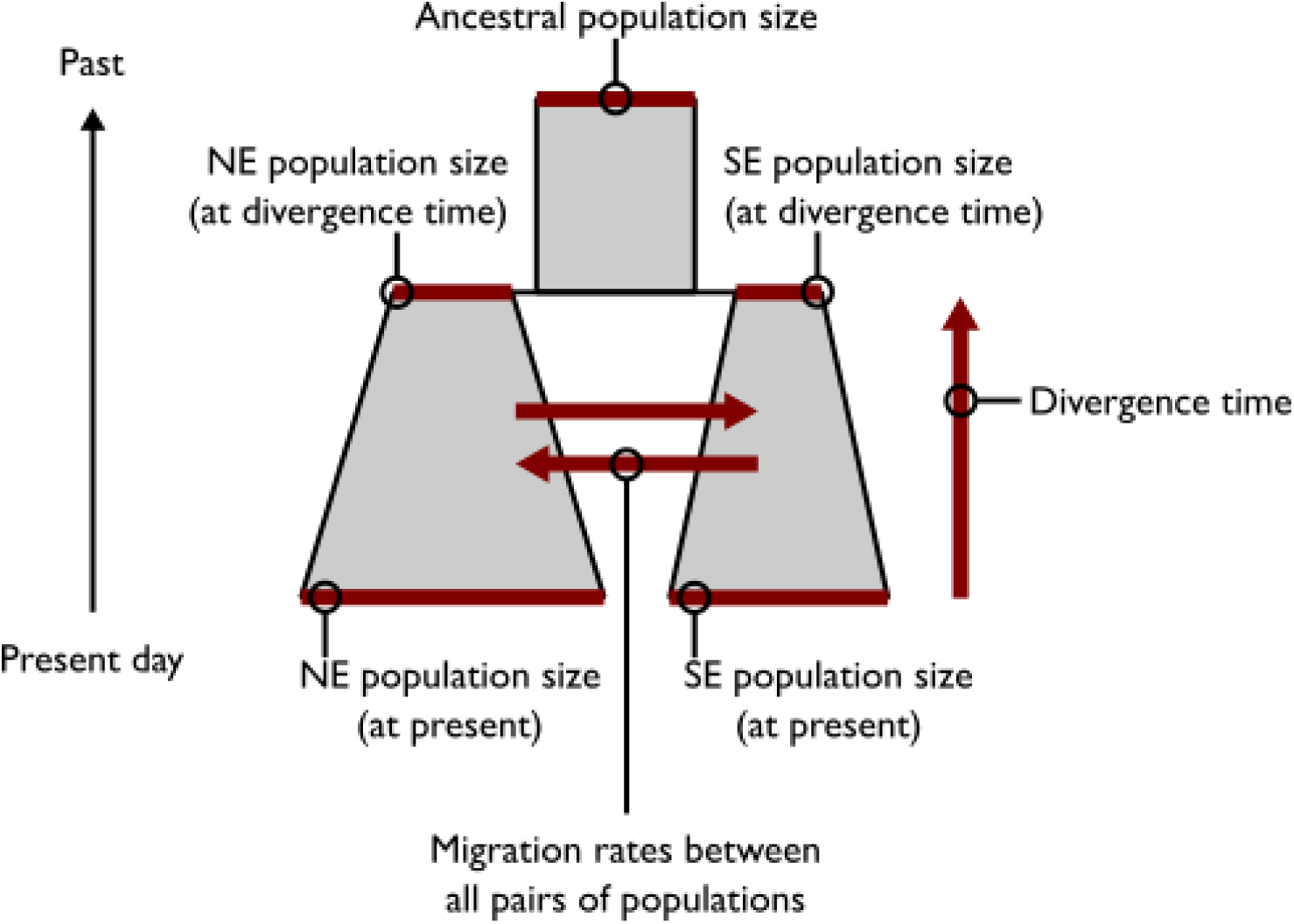
Populations parameters estimated by fastsimcoal2, for *C. mimosae*. Estimated parameters are shown in red. Population growth rates were calculated as log(Population size at divergence time/Population size at present)/Divergence time. Growth rates could be positive or negative depending on the estimates of population sizes. For simplicity, we assumed that the ancestral population(s) were static, without migration between them.

*Results of the coalescent analysis*

Population parameters estimated by fastsimcoal2 coalescent simulations revealed different dynamics in the various species (presented in Tables S5.1 and S5.2). In this analysis, which takes into account population growth and migration, population sizes in the *V. drepanolobium* tree and the ant *C. mimosae* were large relative to *C. nigriceps* and *T. penzigi*, more so than the previously-determined θ values would suggest. We also do not observe that southern population of *C. nigriceps* and *T. penzigi* are particularly large, as suggested by θ.

In addition, we find that the populations of different species at each site are not increasing or decreasing together. For instance, in the highlands sites, *V. drepanolobium* and *C. nigriceps* populations are declining in the northern highlands, but expanding in the southern highlands location, while *C. mimosae* populations are expanding in both, and *T. penzigi* populations are both declining. It is unsurprising that *C. nigriceps* thrives in the Southern Highlands sites, because further to the south and east, some sites have only *C. nigriceps* (Stapley 1999), suggesting that southern Kenya and Tanzania may form a source population for *C. nigriceps*.

Divergence times vary among the different species, with *C. mimosae* having a noticeably shorter divergence time than the other ant species. This may because *C. mimosae* has expanded more recently, or it may be that the ant species expanded together, but that fewer generations of *C. mimosae* have passed in the intervening time. This may well be the case if *C. mimosae* is slower to reproduce, which may likely be true at Mpala at least, given its position on the colonization-competition hierarchy (Stanton *et al.* 2002). Further work on the generation time of the plant and ants will be necessary for these numbers to be interpreted accurately.

Finally, comparing the likelihood of each species’ data under the characteristic population structure of each other species showed some interesting results. In general, the population structure of each species is distinct: forcing the population parameters of any species on any other species resulted in a substantial decrease in likelihood (Table S4.3). There were, however, a few exceptions to this rule. In four cases, there was a small decrease in log-likelihood (less than 1), or even an increase in likelihood by forcing the population structure of on species on another. Most of these involved *C. mimosae*: either forcing the population structure of *C. mimosae* on *V. drepanolobium* or *T. penzigi*, or forcing the population structure of *C. nigriceps* on *C. mimosae*. The fourth case was when the parameters for *V. drepanolobium* were used when estimating *C. nigriceps* data. Since *C. mimosae* is only found in the Northern and Southern Highlands region, these results suggest that much of the differences in population structure among these species are driven by differences in the population sizes of and migration rates to/from the Rift populations.

**Table S4.1:** Population parameter estimates from fastsimcoal2

| Species | Population | Population size (in thousands) |  | Growth rate (forward in time) |  | Divergence time (10<sup>3</sup> generations) |  |
| --- | --- | --- | --- | --- | --- | --- | --- |
|  |  | Mean | Best | Mean | Best | Mean | Best |
| *V. drepanolobium* | NE | 17±1 | 2 | -(1.9±0.2)*10<sup>-3</sup> | -6.7*10<sup>-3</sup> | 43±3 | 30 |
|  | SE | 171±46 | 12 | (8.7±3.0)*10<sup>-4</sup> | -5.2*10<sup>-5</sup> |  |  |
|  | NW | 10±0.4 | 6 | -(6.4±1.6)*10<sup>-5</sup> | 1.4*10<sup>-4</sup> |  |  |
| *C. mimosae* | NE | 71±1 | 62 | (6.4±0.3)*10<sup>-5</sup> | 2.1*10<sup>-5</sup> | 39±1 | 15 |
|  | SE | 108±2 | 139 | (1.6±0.03)*10<sup>-4</sup> | 2.0*10<sup>-4</sup> |  |  |
| *C. nigriceps* | NE | 1.2±0.1 | 1.2 | -(4.3±0.2)*10<sup>-4</sup> | -4.4*10<sup>-4</sup> | 126±3 | 127 |
|  | SE | 3.5±0.4 | 3.4 | (9.3±0.3)*10<sup>-4</sup> | 9.4*10<sup>-4</sup> |  |  |
|  | NW | 4.3±0.1 | 4.3 | -(1.6±0.2)*10<sup>-5</sup> | -1.6*10<sup>-5</sup> |  |  |
|  | SW | 2.1±0.1 | 2.0 | -(2.3±0.1)*10<sup>-3</sup> | -2.3*10<sup>-3</sup> |  |  |
| *T. penzigi* | NE | 1.8±0.02 | 1.7 | -(1.1±0.1)*10<sup>-4</sup> | -6.1*10<sup>-5</sup> | 76±5 | 42 |
|  | SE | 1.9±0.1 | 1.6 | -(7.5±0.8)*10<sup>-4</sup> | -3.1*10<sup>-4</sup> |  |  |
|  | NW | 0.5±0.1 | 0.5 | -(1.5±0.1)*10<sup>-4</sup> | -2.5*10<sup>-4</sup> |  |  |
|  | SW | 1.4±0.01 | 0.9 | -(8.7±0.4)*10<sup>-4</sup> | -1.4*10<sup>-3</sup> |  |  |

Mean values for each parameter are the mean ± standard error, weighted by the likelihood for that run. Divergence time is the number of generations backwards in time until all populations have coalesced.

**Table S4.2:**
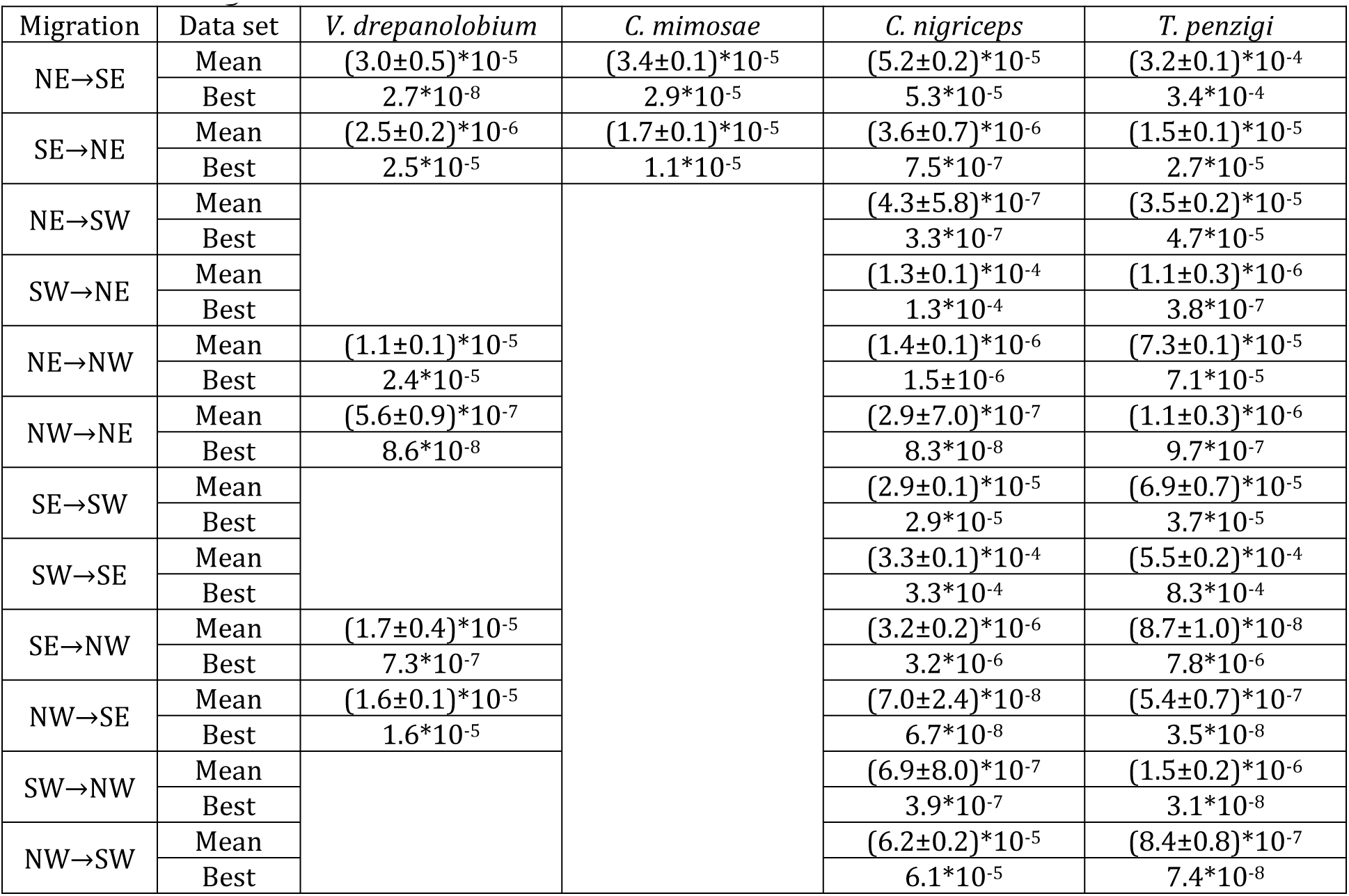
Migration rate estimates from fastsimcoal2

**Table S4.3:**
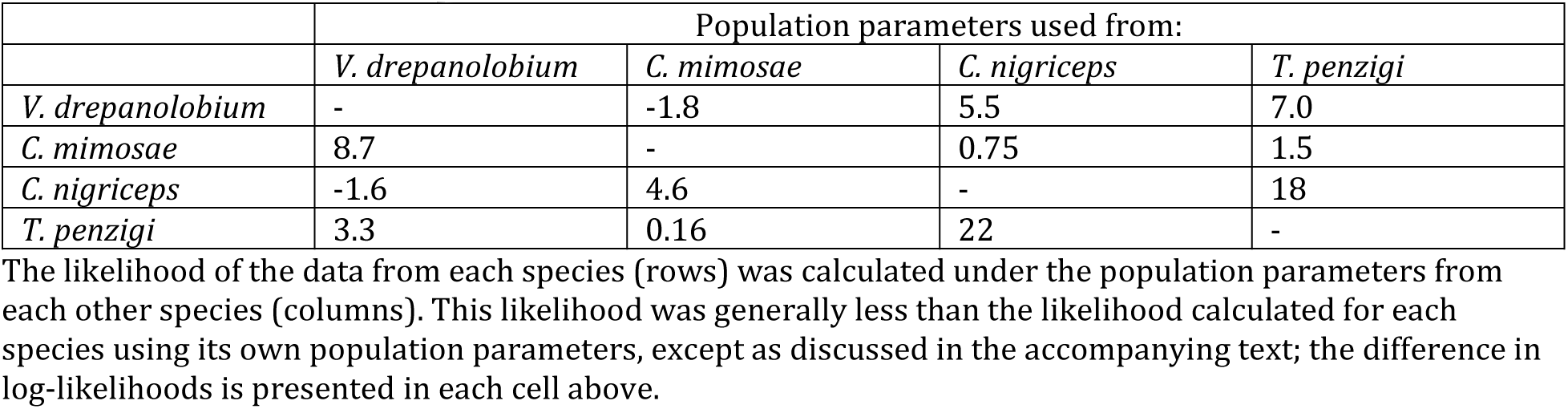
Population genetic data from each species was simulated under the population structure for each other species.

|  | Population parameters used from: |  |  |  |
| --- | --- | --- | --- | --- |
|  | *V. drepanolobium* | *C. mimosae* | *C. nigriceps* | *T. penzigi* |
| *V. drepanolobium* | - | -1.8 | 5.5 | 7.0 |
| *C. mimosae* | 8.7 | - | 0.75 | 1.5 |
| *C. nigriceps* | -1.6 | 4.6 | - | 18 |
| *T. penzigi* | 3.3 | 0.16 | 22 | - |

The likelihood of the data from each species (rows) was calculated under the population parameters from each other species (columns). This likelihood was generally less than the likelihood calculated for each species using its own population parameters, except as discussed in the accompanying text; the difference in log-likelihoods is presented in each cell above.

